# Design and Biophysical Characterization of Second-Generation Cyclic Peptide LAG-3 Inhibitors for Cancer Immunotherapy

**DOI:** 10.1101/2024.08.04.606540

**Authors:** Laura Calvo-Barreiro, Longfei Zhang, Yaser Ali, Ashfaq Ur Rehman, Moustafa Gabr

## Abstract

Lymphocyte activation gene 3 (LAG-3) is an inhibitory immune checkpoint crucial for suppressing the immune response against cancer. Blocking LAG-3 interactions enables T cells to recover their cytotoxic capabilities and diminishes the immunosuppressive effects of regulatory T cells. A cyclic peptide (Cys-Val-Pro-Met-Thr-Tyr-Arg-Ala-Cys, disulfide bridge: 1-9) was recently reported as a LAG-3 inhibitor. Based on this peptide, we designed 19 derivatives by substituting tyrosine residue to maximize LAG-3 inhibition. Screening via TR-FRET assay identified 8 outperforming derivatives, with cyclic peptides 12 [Tyr6(L-3-CN-Phe)], 13 [Tyr6(L-4-NH_2_-Phe)], and 17 [Tyr6(L-3,5-DiF-Phe)] as top candidates. Cyclic peptide 12 exhibited the highest inhibition (IC_50_ = 4.45 ± 1.36 µM). MST analysis showed cyclic peptides 12 and 13 bound LAG-3 with *K*_*D*_ values of 2.66 ± 2.06 µM and 1.81 ± 1.42 µM, respectively, surpassing the original peptide (9.94 ± 4.13 µM). Docking simulations indicated enhanced binding for cyclic peptide 12, with a docking score of -7.236 kcal/mol compared to -5.236 kcal/mol for the original peptide.

---

In 2022, the lymphocyte-activation gene 3 (LAG-3) was approved by the FDA (United States Food and Drug Administration) as the latest immune checkpoint target,^1^ bringing the total number of FDA-approved immune checkpoint inhibitor drugs to four.^2^ However, this anti-LAG-3 cancer immunotherapy, as other monoclonal antibodies before, presents some drawbacks compared to other families of cancer therapies, such as small molecules and peptides. Briefly, small molecules and peptides present more efficient tumor penetration properties, less adverse immune responses over time, lower manufacturing costs, and can be optimized to improve their pharmacokinetic properties.^3-5^ To date, over twenty different peptides are currently used in cancer therapy.^6,7^

Peptides are excellent candidates for targeting protein-protein interactions due to their intrinsic properties, which mimic essential features of proteins, providing them with high specificity.^8^ Consequently, their potential as immune checkpoint inhibitors is being increasingly explored. Chemical modifications are commonly employed to candidate peptides to improve therapeutic properties such as three-dimensional stability, blood circulation time, or shelf-life, among others. One such modification is peptide cyclization. Compared to linear peptides, cyclic peptides offer several advantages in targeting immune checkpoints for cancer immunotherapy. Cyclic peptides are more resistant to enzymatic degradation resulting in extended three-dimensional stability and half-life, thereby improving efficacy.^9^ Therefore, cyclic peptides often show enhanced bioavailability and maintenance of therapeutic levels in the system. The cyclic structure also confers peptides with a conformational rigidity, which preserves the peptide’s 3D conformation and therapeutic properties, avoiding reduced binding affinity or off-target effects. Overall, these characteristics make cyclic peptides promising candidates for cancer immunotherapy, particularly in targeting immune checkpoints where precise modulation of immune responses is crucial for therapeutic efficacy.

Recently, a cyclic peptide targeting LAG-3 immune checkpoint has been discovered using biopanning as the affinity selection technique.^10^ This peptide successfully inhibited LAG-3 interaction with one of its natural ligands, major histocompatibility complex class II (MHC-II).^10^ Based on this peptide’s structure (Cys-Val-Pro-Met-Thr-Tyr-Arg-Ala-Cys, disulfide bridge: 1-9) with submicromolar affinity to LAG-3 protein, we designed 19 different derivatives by incorporating diverse functional groups to the tyrosine amino acid residue (Supplementary Information, Table S1). The conformation of cyclic peptides can be significantly influenced by the chemical composition of the amino acid side chains. Thus, substituting tyrosine with other residues can help stabilize a desired conformation or induce specific structural changes that enhance LAG-3 binding affinity and LAG-3 inhibitory profile.

We first screened the 19 derivatives along with the original cyclic peptide for their ability to inhibit LAG-3/MHC-II interaction using Time Resolved Förster’s Resonance Energy Transfer (TR-FRET) assay.^11^ Briefly, both LAG-3 and MHC-II are tagged with donor and acceptor fluorophores, and LAG-3/MHC-II inhibitors are identified by the reduction in the LAG-3/MHC-II TR-FRET signal. A primary screening using 20 µM as the tested concentration revealed that eight out of the 19 derivatives outperformed the original peptide, being derivatives 12 [Tyr6(L-3-CN-Phe)],13 [Tyr6(L-4-NH_2_-Phe)] and 17 [Tyr6(L-3,5-DiF-Phe)] the three top candidates (Fig. 1A). Subsequent dose-response experiments indicated that cyclic peptide 12 held the highest inhibition capability out of the top candidates, IC_50_ = 4.45 ± 1.36 µM, compared to cyclic peptide 13, IC_50_ = 131.65 ± 35.30 µM; and cyclic peptide 17, IC_50_ = 74.43 µM (Fig. 1B). Regarding cyclic peptide 17, no standard deviation was obtained since two out of the three replicates displayed a wider than accepted IC_50_’s confidence interval and the nonlinear regression model did not adjust the experimental data.

**Fig. 1.**
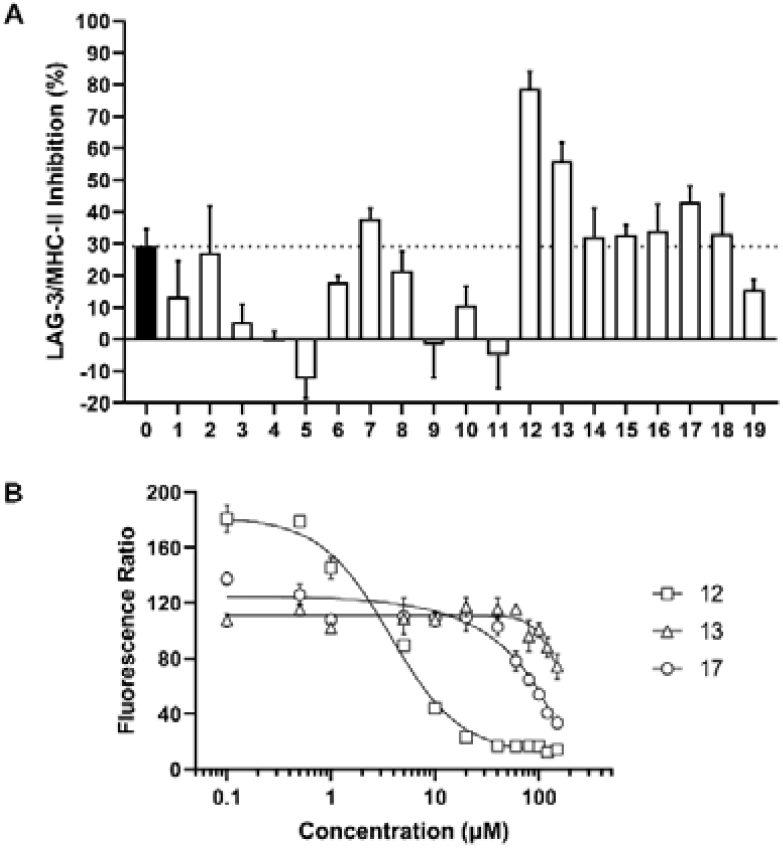
Inhibition of LAG-3/MHC-II interaction by cyclic peptide derivatives. **(A)** Single dosage screening of 19 cyclic peptide derivatives. Tested cyclic peptides at 20 µM (1.5% DMSO) were incubated with Tag1-LAG-3, Tag2-MHC-II, Anti-Tag1 Eu Cryptate reagent, and Anti-Tag2 d2 antibody to study the inhibition ability towards LAG-3/MHC-II interaction. The dashed line indicates the inhibition rate of the original cyclic peptide (0). **(B)** Dose-response curves of the three cyclic peptides in the LAG-3/MHC-II TR-FRET assay. Inhibition rates were measured in triplicate, with results given as the mean ± standard deviation.

We assessed the binding affinity of the original cyclic peptides as well as the three top candidates (cyclic peptide 12, 13 and 17) to LAG-3 protein using microscale thermophoresis (MST) platform, which we validated for affinity screening using fibrinogen-like protein 1 (FGL-1) as a positive control. The binding affinity analysis revealed that cyclic peptides 12 and 13 bound LAG-3 with an equilibrium dissociation constant (*K*_*D*_) equal to 2.66 ± 2.06 µM and 1.81 ± 1.42 µM, respectively (Fig. 2B, C). Both results showed a better performance of these two derivatives compared to the original cyclic peptide: 9.94 ± 4.13 µM (Fig. 2A). Although the screening outcome for cyclic peptide 12 revealed an agreement between LAG-3 binding and inhibition profiles, the outcome for cyclic peptide 13 revealed that protein binding affinity for a ligand does not necessarily correlate with its inhibitory potency.^12^ On the other hand, no *K*_*D*_ value was obtained for cyclic peptide 17. Results regarding cyclic peptide 17 inhibitory capability are in line with LAG-3 binding affinity outcome since no stable IC_50_ values together with no binding to the target protein might indicate unstable binding or false positive results in TR-FRET assays (Fig. 1B).

**Fig. 2.**
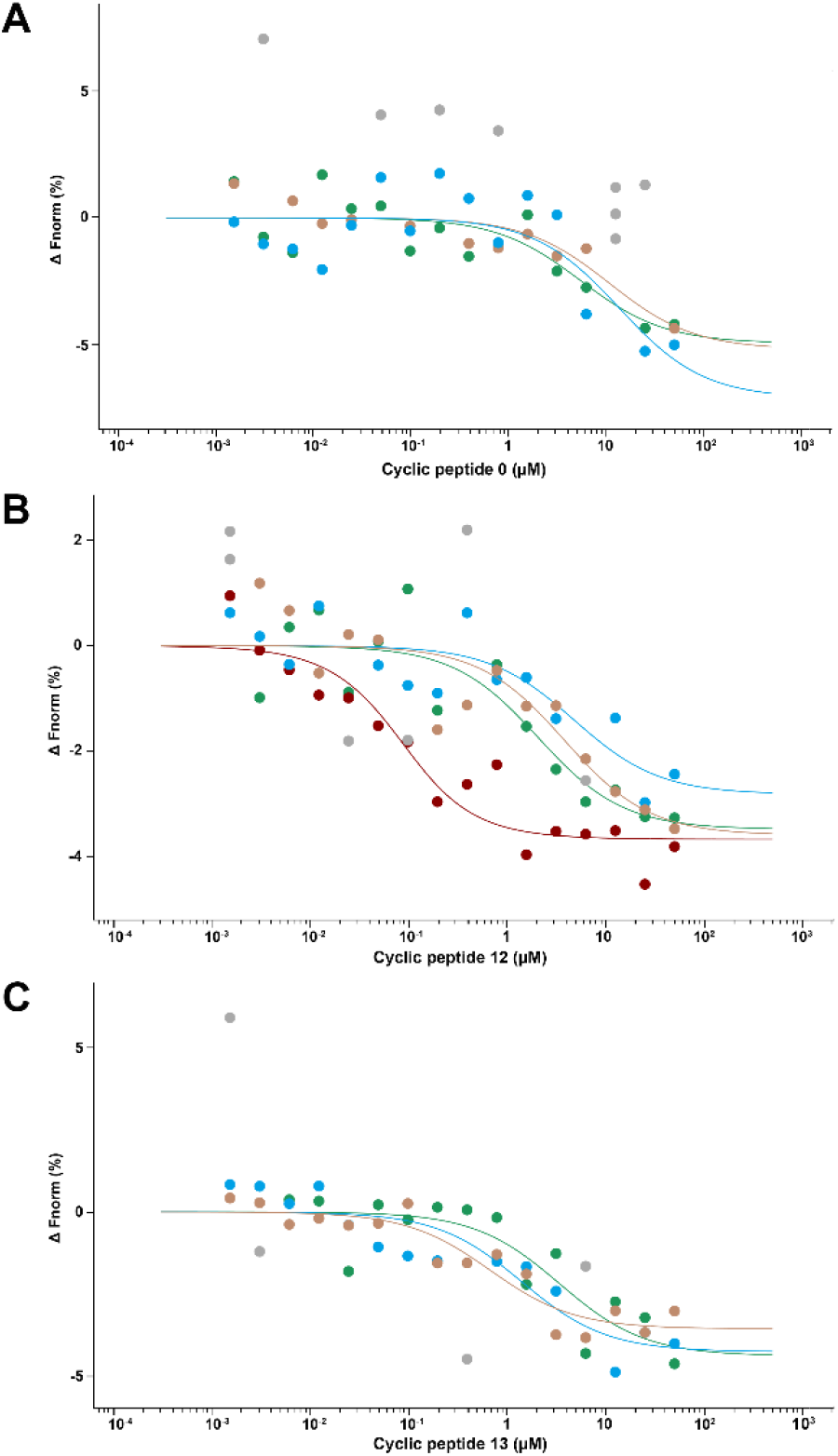
Binding affinity of cyclic peptides 0, 12 and 13 to LAG-3 protein. A range of concentrations (50 µM to 1.53 nM) of the **(A)** cyclic peptid**e** 0, **(B)** cyclic peptide 12 and **(C)** cyclic peptide 13 were incubated with His-labelled human LAG-3 protein (50 nM) for 10 minutes to determine the affinity of the candidate binder for the His-labeled LAG-3 protein using MST. The graphs present the results of three or four independent experiments. Each independent experiment is colored in different colors. Data points in grey color represent outliers/excluded data.

Aiming to investigate the impact of the CN substituent in cyclic peptide 12 in improving LAG-3 binding affinity, we utilized Schrödinger’s GLIDE software for docking simulations. The cyclic peptide 12 not only binds with higher affinity but also aligns more optimally within the complex’s binding pocket compared to the original cyclic peptide (cyclic peptide 0). Specifically, the docking results demonstrated that cyclic peptide 12 achieved an average docking score of -7.236 kcal/mol, an enhancement from -5.236 kcal/mol observed for cyclic peptide 0. The cyclic peptide 12 binds through a two-phase process. Initially, in what we term the tran-conformation (TC), the peptide binds to the pocket but does not completely occupy it, leaving some space unoccupied (Fig. 3A, B). As the peptide relaxes and fully adjusts within the pocket, particularly accommodating the engineered CN group, it transitions to what we refer to as conformation-I (CI). In the TC phase, we observed that the engineered moiety adopts an out-conformation, meaning it protrudes away from the previously unoccupied space, which is now fully occupied due to the peptide’s fully relaxed orientation.

**Fig. 3.**
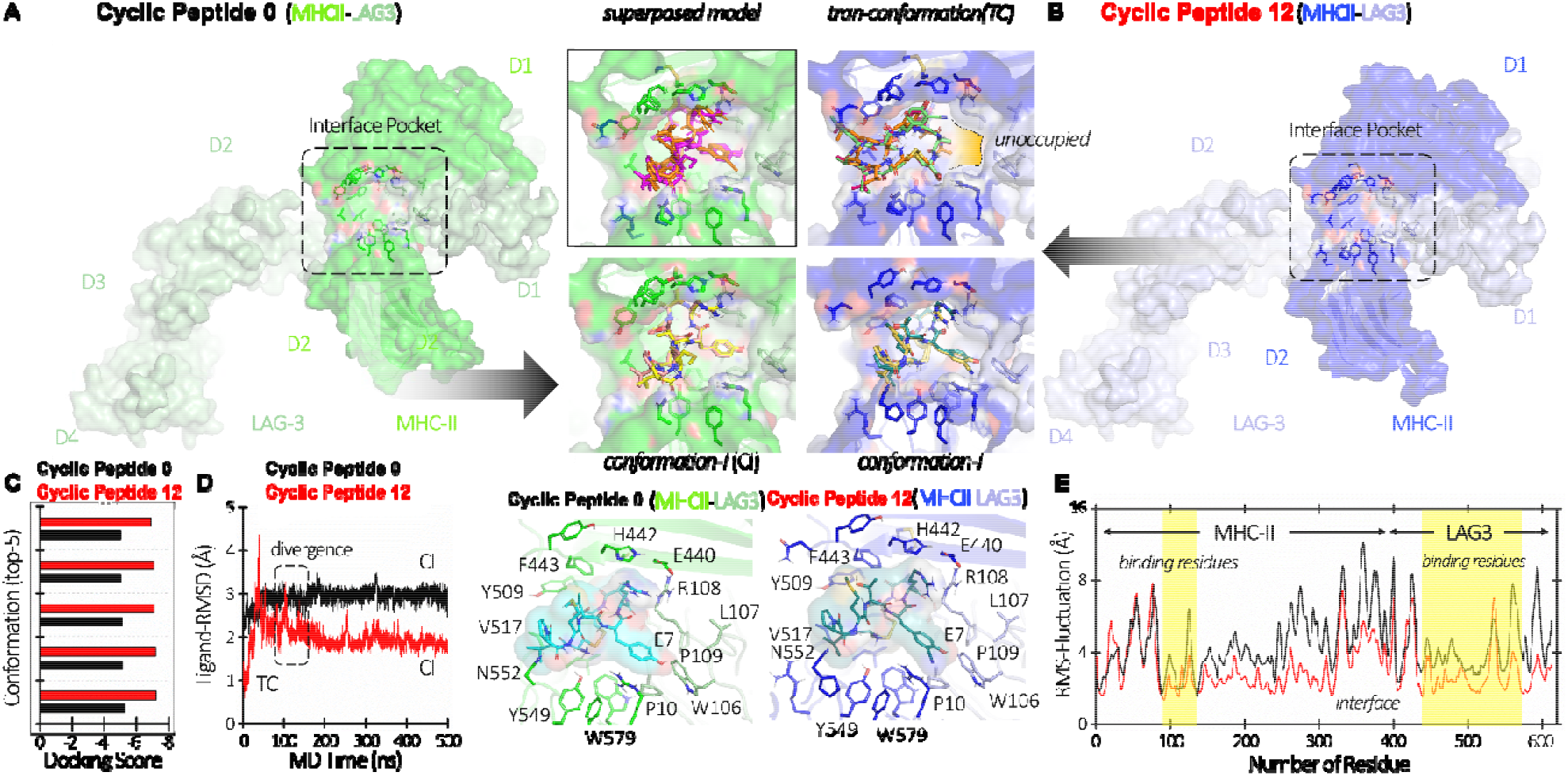
Molecular docking and dynamics analysis of cyclic peptide 0 and cyclic peptide 12 with LAG-3/MHC-II complex. **(A-B)** Cartoon representations of the docked complexes of cyclic peptide 0 and cyclic peptide 12, each domain distinguished by different colors. Zoomed-in sections illustrate the superposition of their top docked conformations (cyclic peptide 0 in pink; cyclic peptide 12 in orange), whit the lower panel illustrating the protein-ligand interaction network, labeling pocket residues. Peptides are rendered in surface representation for clarity. **(C)** Docking scores of th top five docked complexes highlight the impact of CN substitution, increasing the docking score by approximately 38.2%, demonstrating improved binding affinity of cyclic peptide 12 over cyclic peptide 0. **(D)** RMSD profiles demonstrate greater stability of cyclic peptide 12 compared to cycli peptide 0 in the complex. **(E)** RMSF results for individual residues indicate enhanced stability in binding regions, influencing nearby regions allosterically.

The detailed docking analysis revealed that the cyclic peptide 12 binds more effectively due to the CN substitution enhancing π-π interactions and hydrogen bonding within the binding pocket. The CN group facilitates a flipping of the peptide’s 6-ring towards crucial residues W579 on MHC-II and W106 on LAG-3, optimizing π-stacking interactions essential for stable docking. This molecular alignment allows the cyclic peptide 12 to engage more comprehensively with the protein complex, forming multiple hydrogen bonds and hydrophobic interactions with nearby residues including the critical phenylalanine F443 (Fig. 3A, B).

In our simulations, we additionally analyzed the dynamic behavior of cyclic peptide 0 and cyclic peptide 12 when complexed with LAG-3 and MHC-II over a total simulation time of 1.0 µs, with each simulation run lasting 0.5 µs. Notably, c**y**clic peptide 12 exhibited a progressive decrease in root-mean-square deviations (RMSD), suggesting that it systematically explored stable orientations within the pocket by adjusting laterally and interacting with critical interface residues (Fig. 3D). This stability is corroborated by root-mean-square fluctuation (RMSF) analyses, which highlighted decreased flexibility in key binding regions, suggesting a more effective interaction with the target protein complex (Fig. 3E). Moreover, our observations indicated that cyclic peptide 12 initially bound to LAG-3 and MHC-**II** in a manner where it entered the pocket without fully occupying it. However, by the 200 ns mark, we noted the peptide beginning to adjust within the pocket and ultimately adopting a conformation similar to that of cyclic peptide 0, albeit with a more snug fit orientation (Fig. 3D).

The analysis of the top conformations from the docking studies emphasized that cyclic peptide 12, despite slight initial fluctuations, found a stable orientation within the binding pocket faster than cyclic peptide 0 (Fig. 3C, D). These fluctuations did not indicate instability; instead, they represented the peptide’s process of adjusting within the space to achieve optimal fit. This dynamic adjustment process is essential for the prolonged interaction of cyclic peptide 12 with LAG-3, which **w**ould significantly enhance the therapeutic potential of the peptide.

Considering the promising results of cyclic peptide 12 in both biochemical assays and docking simulations, we next eval**u**ated the ability of the derivative peptide to inhibit tumor growth *in vivo*. CT26.WT tumor-bearing mice were treated with 8 mg/kg/day for 21 days, and no differences were observed in either tumor growth or survival rates between the groups (Fig. 4A, B). Since cyclic peptide 0 demonstrated antitumor effects via CD8^+^ T cells, we aimed to investigate the potential immunomodulatory effect of cyclic peptide 12 in the tumor local environment (Supplementary Information, Fig. S1-2). However, no difference was observed for any of the studied immune populations: helper T cells (total, Th1 and Th17), cytotoxic T cells (total and IFN-γ-expressing cells), regulatory T cells, tumor-specific CD4+ T cells, and tumor-specific CD8^+^ T cells (Fig. 4C-J).

**Fig. 4.**
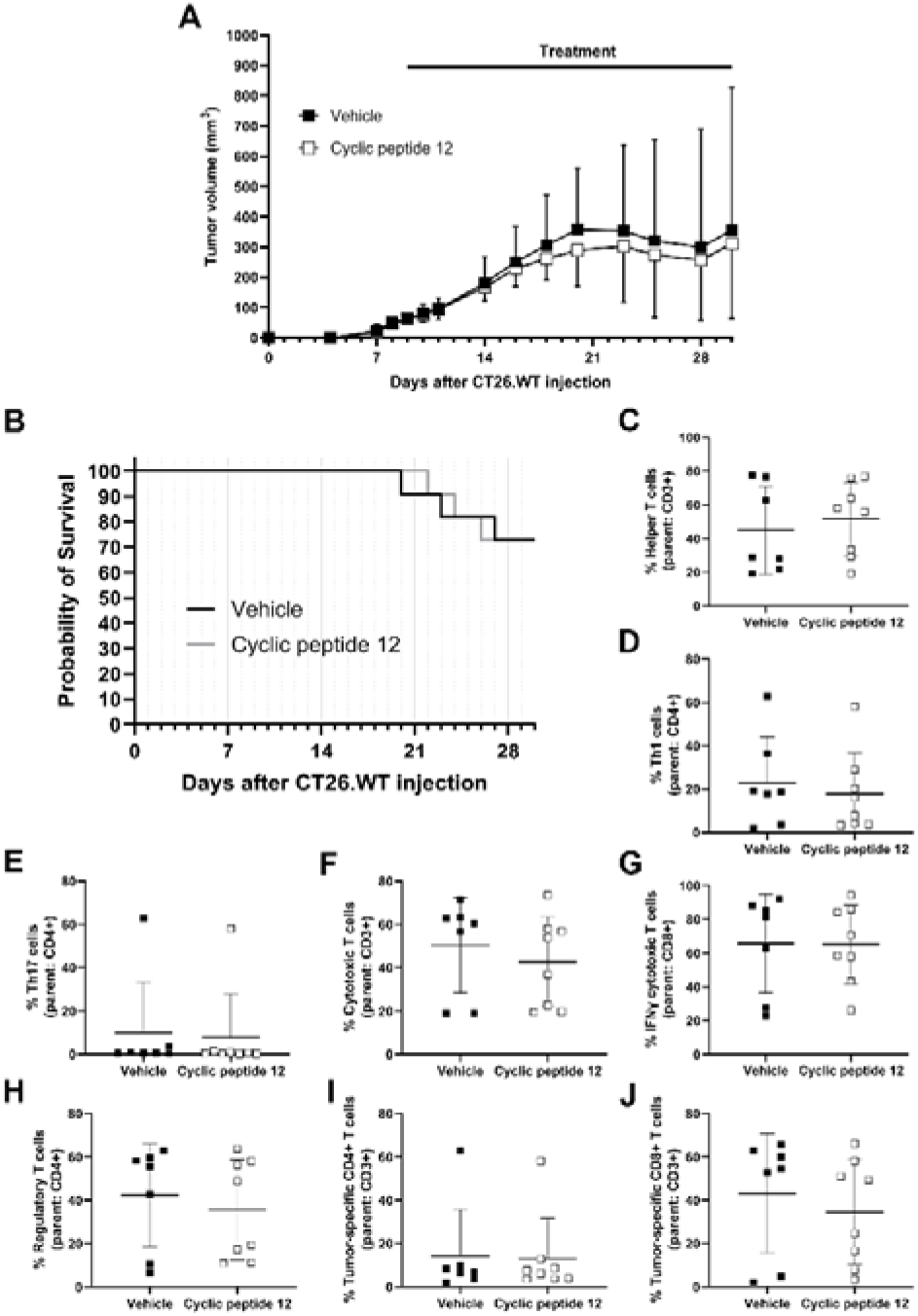
Effect of cyclic peptide 12 therapeutic treatment on tumor growth and tumor infiltrating immune cells in CT26.WT tumor bearing mice. **(A)** CT26.WT tumor bearing mice were treated with 8 mg/kg/day (intraperitoneal administration) for three weeks from day 9 after CT26.WT injection. Treatment of mice was initiated after the tumors had been grown until reaching a palpable size of 40-100 mm^3^. The graph presents one independent experiment (Vehicle, n=11 and Cyclic peptide 12, n=11). The data are presented as the mean ± standard deviation. **(B)** Overall mice survival (in days) after the injection of CT26.WT colon carcinoma cell line (500,000 cells/animal). The graph presents one independent experiment (total number of animals at day 0: Vehicle, n=11 and Cyclic peptide 12, n=11). Percentage of **(C)** helper T cells, **(D)** Th1 cells, **(E)** Th17 cells, **(F)** cytotoxic T cells, **(G)** IFN-γ-expressing cytotoxic T cells, **(H)** regulatory T cells, **(I)** tumor-specific helper T cells, and **(J)** tumor-specific cytotoxic T cells in the tumor microenvironment. The charts present the results of an independent experiment (Vehicle, n=7 and Cyclic peptide 12, n=8). The data are presented as the means ± standard deviations. Abbreviations: Th: Helper T cell.

The absence of immune modulation within the tumor microenvironment led us to speculate whether cyclic peptide 12 might have lost its structural stability upon injection into the animals. To investigate this, we conducted peptide stability assays using both cyclic peptide 0 and cyclic peptide 12 in mouse and human serum over a 24-hour period (Fig. S43). Cyclic peptide 0 remained intact after 24 hours of incubation in mouse serum (Fig. S43A), and similar results were observed in human serum (Fig. S43B). Similarly, the cyclic peptide 12 structure was not affected by mouse or human serum (Fig. S43C, D). The single peak observed for both cyclic peptides in each experimental condition indicates their stability in both types of serum. However, both cyclic peptides appeared to non-specifically bind to mouse and human serum proteins, resulting in a reduced abundance over time, as evidenced by the filtration process prior to HPLC analysis (Fig. S43). Something similar might have been occurring in vivo and so, no effective quantities would have reached LAG-3-expressing immune cells, which could potentially explain the lack of observed effectiveness.

Conversely, our results are consistent with previous studies demonstrating a significant presence of PD-1-expressing CD8+ T cells within tumor infiltrating lymphocytes (TILs) in CT26 tumor-bearing mice (Fig. 4J).^13^ PD-1, another negative immune checkpoint, is known to suppress anti-tumor immune responses. Consistent with observations in other tumor models,^14-16^ our data on LAG-3 monotherapy inefficacy underscore the necessity of combining anti-LAG-3 with anti-PD-1/PD-L1 blockade. Such combination therapies are crucial not only for reducing tumor growth but also for exerting immunomodulatory effects within TILs.

These results provide compelling evidence of the effectiveness of molecular modifications in cyclic peptides for therapeutic applications. The introduction of a CN substituent not only optimizes therapeutic interactions but also significantly enhances the binding efficiency and stability of cyclic peptides, offering a robust platform for the development of potent immunotherapeutic agents targeting the LAG-3/MHC-II interaction. This study underscores the critical role of detailed molecular understanding in the design and optimization of peptide-based therapeutics and highlights the importance of dual immunotherapies to achieve therapeutic effects.

## Data Availability

Data will be made available on request.

## Acknowledgments

The authors thank Dr. Natalie Fuchs for technical assistance on processing tumor samples for subsequent flow cytometry analysis. Microscale Thermophoresis (MST) was performed in the Rockefeller University’s Bio-Imaging Resource Center, RRID:SCR_017791. We gratefully acknowledge financial support from the ELSA U. Pardee Foundation (Award ID: 2022-215).

