## Supporting Information for "Design and Biophysical Characterization of Second-Generation Cyclic Peptide LAG-3 Inhibitors for Cancer Immunotherapy"

**5. Stability of the peptides in human and mouse serum…………….………………………………..……………………32**

### 1. Materials & Methods

#### 1.1. Synthesis of peptides

The peptides were synthesized on 2-CL resin, using standard Fmoc synthesis protocol with DIC/Cl-HOBt coupling, on an APEX 396 automatic synthesizer.  The resin was swollen in DMF for 30 min, treated with 20v% Piperidine-DMF for 8 minutes at 50°C to remove the Fmoc protecting group and washed with DMF for three times.  For the coupling reaction, the resin was added with Fmoc-protected amino acid, Cl-HOBt, DIC and NMP.  The mixture was vortexed for 20 minutes at 50°C.  Afterwards, the resin was washed with DMF once.  The cycle of deprotection and coupling steps was repeated until the last amino acid residue was assembled.  After the final Fmoc protecting group was removed, the resin was treated with 20v% acetic Anhydride-NMP for 20 minutes.  The resin was then washed with DMF, DCM and dried with air. The peptides were cleaved using a TFA cocktail (95v%TFA, 2.5v%water and 2.5v%TIS) for three hours.  Crude peptides were precipitated by adding ice-chilled anhydrous ethyl ether, washed with anhydrous ethyl ether for three times, and freeze-dried. Air oxidation reactions were carried out on the freeze-dried crude peptides, which were dissolved in water-acetonitrile (pH < 7, adjusted with (NH4)2CO3), to form the disulfide bond. After disulfide cyclization, the crude peptides were loaded onto a prep-HPLC column and purified with a gradient of 10%-55%B within 45 minutes at a flow rate of 12 ml/min.  The product was analyzed by LC-MS and confirmed to have >95% HPLC purity, and freeze-dried.

#### 1.2. Time-Resolved Förster’s Resonance Energy Transfer (TR-FRET) Assay

TR-FRET assay was performed using a commercially available kit (Cat #64LAG3PEG, Cisbio, Waltham, MA, USA). Briefly, in a 96-well low-volume white assay plate, 2 µL tested peptide was mixed with 4 µL Tag1-LAG-3, 4 µL Tag2-MHC-II, 5 µL Anti-Tag1 Eu Cryptate reagent, and 5 µL Anti-Tag2 d2 antibody, and the final concentrations of peptides were 150, 120, 100, 80, 60, 40, 20, 10, 5, 1, 0.5, and 0.1 µM (n=3). The mixture of Tag1-LAG-3, Tag2-MHC-II, Anti-Tag1 Eu Cryptate reagent, and Anti-Tag2 d2 antibody was used as the positive control, and the mixture without Tag2-MHC-II was used as the negative control. After incubating at room temperature for 3 h with gentle shaking, fluorescence intensities were acquired on a Tecan Infinite M1000 Pro equipment (λ_ex_: 340 nm, λ_em_: 620 nm and 665 nm; Tecan, Männedorf, Switzerland). The signals were calculated as a ratio: Ratio = Fluorescence_665 nm_ / Fluorescence_620 nm_ × 10^4^. All assay wells contained 1.5% DMSO.

#### 1.3. Microscale Thermophoresis (MST)

The Protein Labeling Kit RED-NHS 2^nd^ Generation (Cat #MO-L011, NanoTemper Technologies, Munich, Germany) was used for the labeling of the human LAG-3 (hLAG-3) protein, His tag (Cat # 16498-H08H, Sino Biological, Beijing, China) following manufacturers’ instructions. No buffer exchange was needed before protein labeling and purification. Briefly, a 4.4 µM hLAG-3 protein solution was mixed with a 5.5-fold concentration of RED-NHS 2^nd^ Generation dye in assay buffer and incubated for 30 minutes at room temperature in the dark. The assay buffer was 1x PBS, 0.05% Tween 20, pH 7.4. After protein-dye incubation, LAG-3 purification by removing the remaining free dye was performed following manufacturers’ instructions. The final concentration and the degree of labeling (DOL) of hLAG-3, the number of dye molecules attached to individual protein molecules, were calculated based on absorbance measurements (A_280_ and A_650_, respectively) using a Biotek Synergy Neo2 Reader (Agilent, Santa Clara, CA, USA). Successful labeling was considered if the DOL was comprised between 0.5 and 1. Additionally, the protein fold was confirmed by comparing the melting curve of hLAG-3 before and after protein labeling using the Tycho NT.6 system (NanoTemper Technologies). Finally, labeled hLAG-3 protein was flash-frozen in liquid nitrogen and stored at -80°C until use.

Compounds were diluted in MST buffer (10 mM HEPES, 150 mM NaCl, 0.005% Tween 20, pH 7.4) to their respective final concentrations in the presence of 2% DMSO. Human fibrinogen-like protein 1 (FGL-1), Fc tag (Cat #FG1-H5258, Acro Biosystems) in MST buffer supplemented with 2% DMSO served as positive controls, while MST buffer with 2% DMSO alone served as the negative control.

Before each use, freshly thawed labeled LAG-3 stock protein underwent centrifugation at 15,000 rpm and 4°C for 10 minutes. Subsequently, the protein was diluted to 100 nM in MST buffer and incubated in a 1:1 ratio with the corresponding 2-fold concentrated cyclic peptide or control, resulting in a final volume of 20 µL. This mixture was incubated for 10 minutes at room temperature in the dark.

All measurements were conducted using a NanoTemper Monolith NT.115 instrument (NanoTemper Technologies) with Monolith Standard Capillaries (Cat #MO-K022, NanoTemper Technologies). The system parameters were set to 25 °C, 40% excitation power and medium MST power, and fluorescence was measured. Initial measurements were taken for 1 second without heating (prior to infrared laser activation), followed by measurements during 21 seconds with the infrared laser activated. Results were recorded as follows: i) relative fluorescence, which represents fluorescence during the experiment duration (TRIC) normalized to initial fluorescence at the set temperature; ii) normalized fluorescence (Fnorm, %), calculated as the ratio of fluorescence values after laser activation to those before.

For each selected cyclic peptide, a 16-point serial dilution was prepared starting at 50 µM. Affinity constants were determined based on three to four independent experiments.

#### 1.4. *In Vitro* Stability in Serum

56.2 µL of cyclic peptide (20 mM, PBS) was added into 505.8 µL mouse (Cat #10410, Invitrogen, Waltham, MA, USA) or human serum (Cat #H4522, Sigma-Aldrich), and the mixtures were incubated at 37 °C. At selected time points (0, 2, 4, 6, 8, 11, and 24 h), a 50 µL sample was collected and treated with 150 µL of acetonitrile (ACN). After centrifugation and filtration, the supernatant was collected and subjected to high-performance liquid chromatography (HPLC; elution A: water with 0.1% TFA, elution B: ACN with 0.1% TFA) using gradient elution from 0 to 95% of elution B at a flow rate of 0.5 mL/min. The data were quantified by analyzing peak areas, and the residual ratios were expressed as a percentage of the initial content.

#### 1.5. Computational Study

The LAG-3/MHC-II complex (from our previous study^1^) was prepared using the Maestro protein preparation wizard with default parameters and the OPLS4 forcefield. This was followed by restrained minimization of the protein structure until the continued the average root-mean-square deviation (ARMSD) of heavy atom reached 0.30 Å^2^. The Glide Grid for peptide docking was generated using the cryptic binding site of SA-15 to define the cavity^1^. Physiological pH was used for residue protonation. The wild-type cyclic peptide (Cyclic Peptide 0) and mutant cyclic peptide (Cyclic Peptide 12) models were built using Maestro molecule builder panel.

The ground state electronic structure energy optimization for both WT-cyclicPep and MT-cyclicPep was carried out using hybrid density functional theory (DFT) with the Becke, three-parameter, Lee–Yang–Parr with dispersion correction (B3LYP-D3) wave function and the basis sets of 6-311 + G(d,p) within Schrödinger’s Jaguar software package. Before docking, the default parameters for the conformational search (Mixed Torsional/Low-mode sampling algorithm) were considered, generating 64 and 256 conformers for the wild-type and mutant peptides, respectively. All the generated conformers were used as starting point for docking. The SP-Peptide module from Glide 6.5 were used to perform the docking of all possible conformations of both the peptides with default parameters. A total of 2,926 and 8,295 poses were generated from the Glide SP-peptide docking simulations. The best-docked poses were chosen based on Glides docking energy, Glide energy, and Glide Emodel energy. The top five final poses for both peptides were subjected to MMGBSA to further optimize the pose conformation. The OPLS4 force field and GB/SA continuum solvent model were used to validate the accuracy of the docking score, confirming the stability of the docking complex.

Molecular dynamics (MD) simulations of 1 µs were conducted using the Desmond-v7.2 software, integrated with the Maestro suite^3^. The complexes were placed in an orthorhombic box implementing the periodic boundary conditions of 10 Å and filled with explicit water molecules using simple point charge (SPC) water model. The MD calculations utilized the OPLS4 force field^4^. To mimic physiological conditions, the system was neutralized with Na^+^ and Cl^−^ ions using a salt concentration of 0.15 M. Simulations were run in the NPT ensemble, maintaining a constant temperature of 300 K and a pressure of 1.01325 bar. The duration of the simulation was set to 500 ns for each complex, with data collection every 20 ps. The short-range interactions were calculated within a 9.0 Å cut-off radius. Temperature and pressure were regulated using the Nose–Hoover thermostat^5^ and the Martyna–Tobias–Klein^6^ methods, respectively. The equations of motion were integrated using the RESPA integrator with a 2.0 fs time step for both bonded and non-bonded interactions within the cut-off range^7^. Before starting the simulations, the system was subjected to minimization and equilibration processes in accordance with the default protocols provided by Desmond. For analyzing the trajectory, the simulation interaction diagram protocol from the Desmond package was utilized. At the end high resolution images were rendered using PyMol Molecular Graphics system^8^.

#### 1.6. Mice

BALB/cJ (Stock #: 000651) five- to six-week-old female mice purchased from Jackson Laboratory (Farmington, CT, USA) were used. Mice were housed under standard light- and climate-controlled conditions, and standard chow and water were provided *ad libitum*. All mouse experiments complied with ARRIVE (Animal Research: Reporting of In Vivo Experiments) guidelines and were carried out in accordance with the NIH (National Research Council) Guide for the Care and Use of Laboratory Animals and the protocols approved by the Institutional Animal Care and Use Committee (IACUC) at Weill Cornell Medicine (protocol number: 2023-0022).

#### 1.7. Injection and Assessment of CT26 Animal Model

Five- to six-week-old female BALB/c mice were subcutaneously injected with 500,000 syngeneic CT26 cells into the dorsal area to establish colorectal cancer model. Tumor sizes were measured using a digital caliper, and tumor volumes were calculated as: volume (mm^3^) = 0.5 × length (mm) × width (mm) × width (mm). Treatment of mice was initiated after the tumors had been grown for 8 to 10 days until reaching a palpable size of 40-100 mm^3^. At day of randomization, tumor bearing mice were grouped in the two experimental groups (11 mice per group) using individual tumor volume (mm^3^) as classification criteria. The following endpoint criteria were used for mouse euthanasia: i) tumor volume higher than 1.5 cm^3^, ii) ulceration size higher than 20% of tumor size, and iii) animal classified with a body condition score (BCS) equal to 2 (underconditioned mouse) that cannot be sustained beyond 3 days without extensive supportive care: frequent rehydration, special foods, and heat.

#### 1.8. Tumor Collection and Dissociation to Single-Cell Suspensions

At 30 days post injection, tumor-bearing mice were euthanized by carbon dioxide asphyxiation. After pinning down and spraying the animals with 70% ethanol, subcutaneous tumor samples were collected using sterilized tweezers. Skin, fat, and fibrous areas were removed from the tumor sample. All collected tumors were put into a tube containing pre-chilled MACS^®^ tissue storage solution (130-100-008, Miltenyi Biotec, Bergisch Gladbach, Germany) and kept overnight at 4°C.

The Tumor Dissociation Kit (130-096-730, Miltenyi Biotec) was used for dissociation of tumor tissues into single suspensions using previously described protocol^9^.

#### 1.9. Flow Cytometry Analysis

Single-cell suspensions were prepared as described above. Cell subsets were analyzed using fluorochrome-conjugated monoclonal antibodies (mAbs) after discrimination of dead cells by Fixable Viability Stain (BD Pharmingen, BD Biosciences, Franklin Lakes, NJ, USA). For analysis of the helper, cytotoxic and regulatory T cell populations, CD45 (103155, BioLegend, San Diego, CA, USA), CD3ε (553062, BD Pharmingen), CD4 (46-0042, eBioscience, San Diego, CA, USA), CD8a (560182, BD Pharmingen), and CD25 (12-0251, eBioscience) were used. FoxP3 intracellular staining was performed using fluorochrome-labelled anti-FoxP3 mAb (17-5773, eBioscience). PD-1 (135221, BioLegend) was selected and evaluated in the T cell populations for tumor-specific cell classification. For intracellular cytokine determination, *ex vivo* stimulation of splenocytes was performed with 50 ng/ml phorbol 12-myristate 13-acetate (PMA, P1585, Sigma-Aldrich) and 1 µg/ml ionomycin (I0634, Sigma-Aldrich) in the presence of GolgiPlug and GolgiStop (555029 and 554724, BD Biosciences) for 4 h. Then, CD45 (103155, BioLegend), CD3ε (553062, BD Pharmingen), CD4 (46-0042, eBioscience), and CD8a (560182, BD Pharmingen) staining was performed. Cytokine intracellular staining was performed using fluorochrome-labelled anti-IFN-γ (554413, BD Pharmingen), and anti-IL-17A (559502, BD Pharmingen) mAbs. Fluorescence was analyzed with a BD LSRFortessa™ Cell Analyzer (BD Biosciences) and FlowJo 10.10.0 software (Ashland, OR, USA).

### 2. Flow Cytometry Gating Strategy

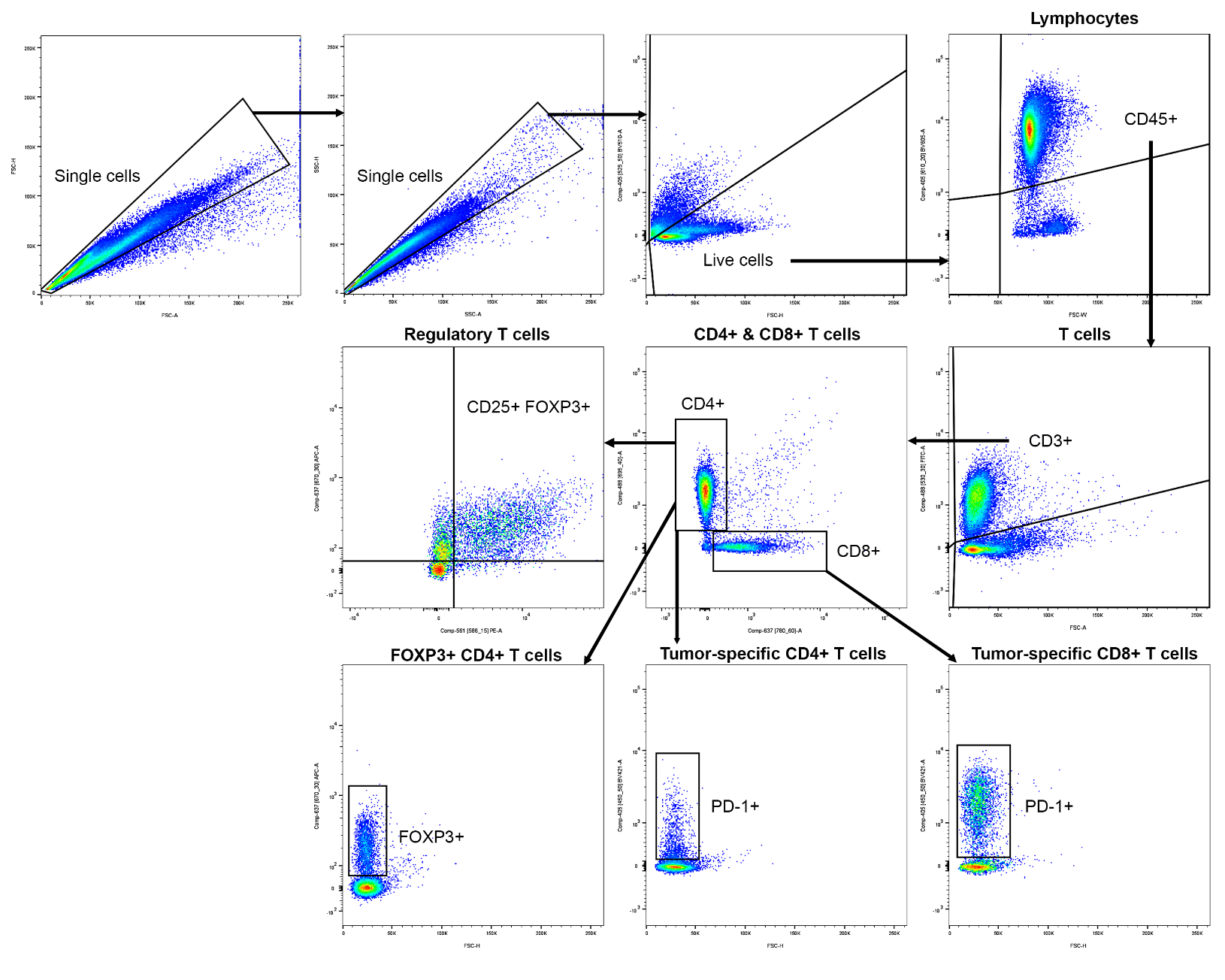

**Figure S1.** Flow cytometry gating strategy for helper, cytotoxic, regulatory, and tumor-specific T cell populations.

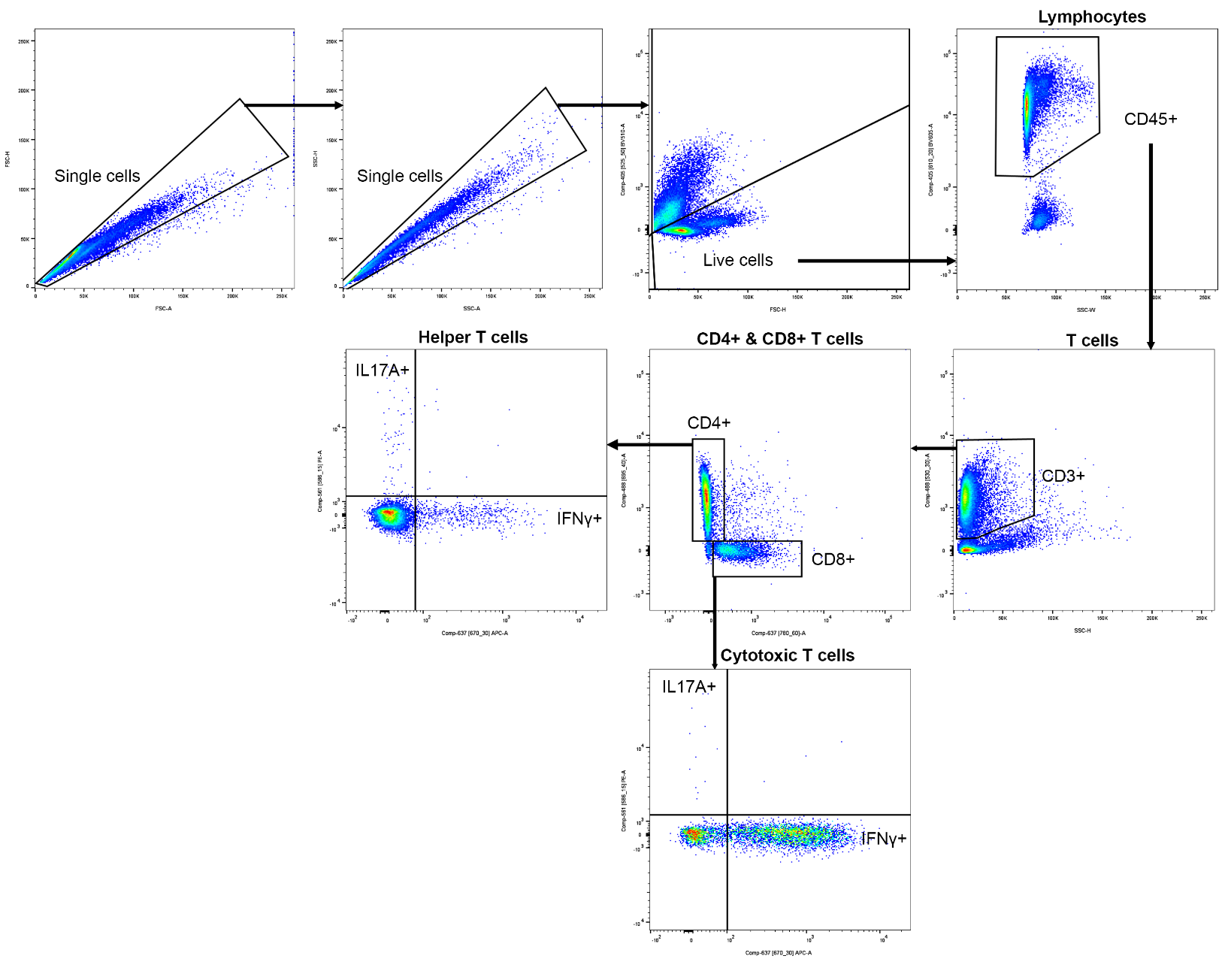

**Figure S2.** Flow cytometry gating strategy for IL-17A- and IFN-γ-producing T cell populations.

### 3. Structures of Cyclic Peptides

**Table S1.** Structures of original cyclic peptide (0) and cyclic peptide derivatives (1 to 19).

| **Peptide ID** | **Sequence** |
| --- | --- |
| Cyclic peptide 0 | Cys-Val-Pro-Met-Thr-**Tyr**-Arg-Ala-Cys |
| Cyclic peptide 1 | Cys-Val-Pro-Met-Thr-**Phe**-Arg-Ala-Cys |
| Cyclic peptide 2 | Cys-Val-Pro-Met-Thr-**L-4-F-Phe**-Arg-Ala-Cys |
| Cyclic peptide 3 | Cys-Val-Pro-Met-Thr-**L-4-Cl-Phe**-Arg-Ala-Cys |
| Cyclic peptide 4 | Cys-Val-Pro-Met-Thr-**L-4-Me-Phe**-Arg-Ala-Cys |
| Cyclic peptide 5 | Cys-Val-Pro-Met-Thr-**L-4-CN-Phe**-Arg-Ala-Cys |
| Cyclic peptide 6 | Cys-Val-Pro-Met-Thr-**L-4-NO2-Phe**-Arg-Ala-Cys |
| Cyclic peptide 7 | Cys-Val-Pro-Met-Thr-**Cha**-Arg-Ala-Cys |
| Cyclic peptide 8 | Cys-Val-Pro-Met-Thr-**L-1-Nal**-Arg-Ala-Cys |
| Cyclic peptide 9 | Cys-Val-Pro-Met-Thr-**L-3-F-Phe**-Arg-Ala-Cys |
| Cyclic peptide 10 | Cys-Val-Pro-Met-Thr-**L-3-Cl-Phe**-Arg-Ala-Cys |
| Cyclic peptide 11 | Cys-Val-Pro-Met-Thr-**L-4-I-Phe**-Arg-Ala-Cys |
| Cyclic peptide 12 | Cys-Val-Pro-Met-Thr-**L-3-CN-Phe**-Arg-Ala-Cys |
| Cyclic peptide 13 | Cys-Val-Pro-Met-Thr-**L-4-NH2-Phe**-Arg-Ala-Cys |
| Cyclic peptide 14 | Cys-Val-Pro-Met-Thr-**L-3-Cl-Tyr**-Arg-Ala-Cys |
| Cyclic peptide 15 | Cys-Val-Pro-Met-Thr-**L-3-I-Tyr**-Arg-Ala-Cys |
| Cyclic peptide 16 | Cys-Val-Pro-Met-Thr-**L-3,5-DiI-Tyr**-Arg-Ala-Cys |
| Cyclic peptide 17 | Cys-Val-Pro-Met-Thr-**L-3,5-DiF-Phe**-Arg-Ala-Cys |
| Cyclic peptide 18 | Cys-Val-Pro-Met-Thr-**L-4-Br-Phe**-Arg-Ala-Cys |
| Cyclic peptide 19 | Cys-Val-Pro-Met-Thr-**L-3-NO2-Tyr**-Arg-Ala-Cys |

### 4. HPLC and MS Data for Cyclic Peptides

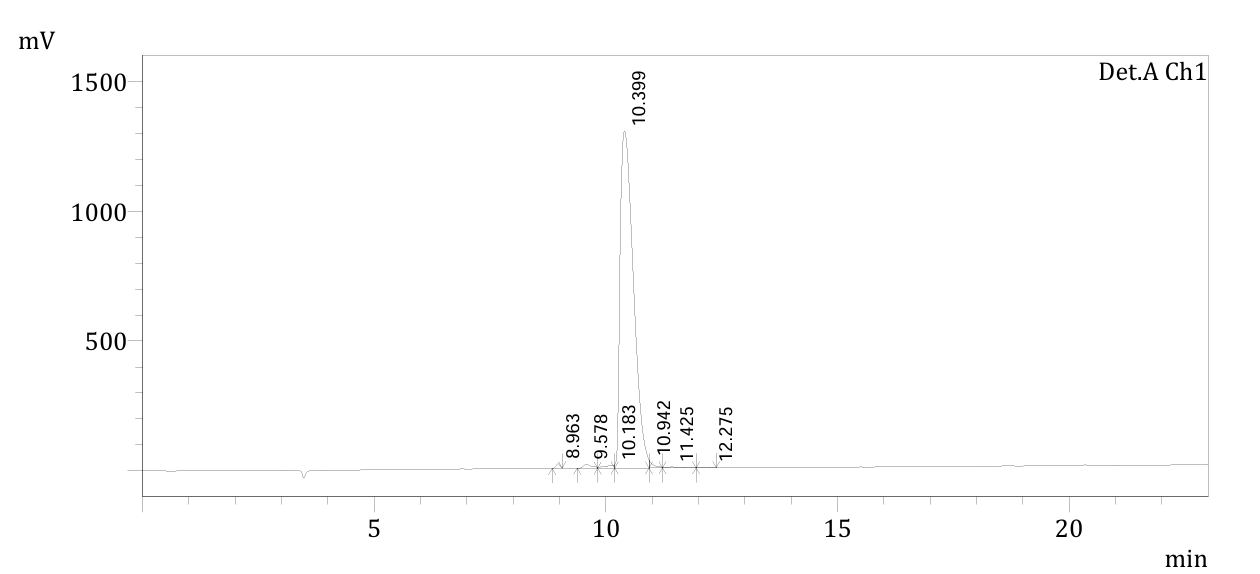

**Figure S3.** HPLC trace for Cyclic peptide 0.

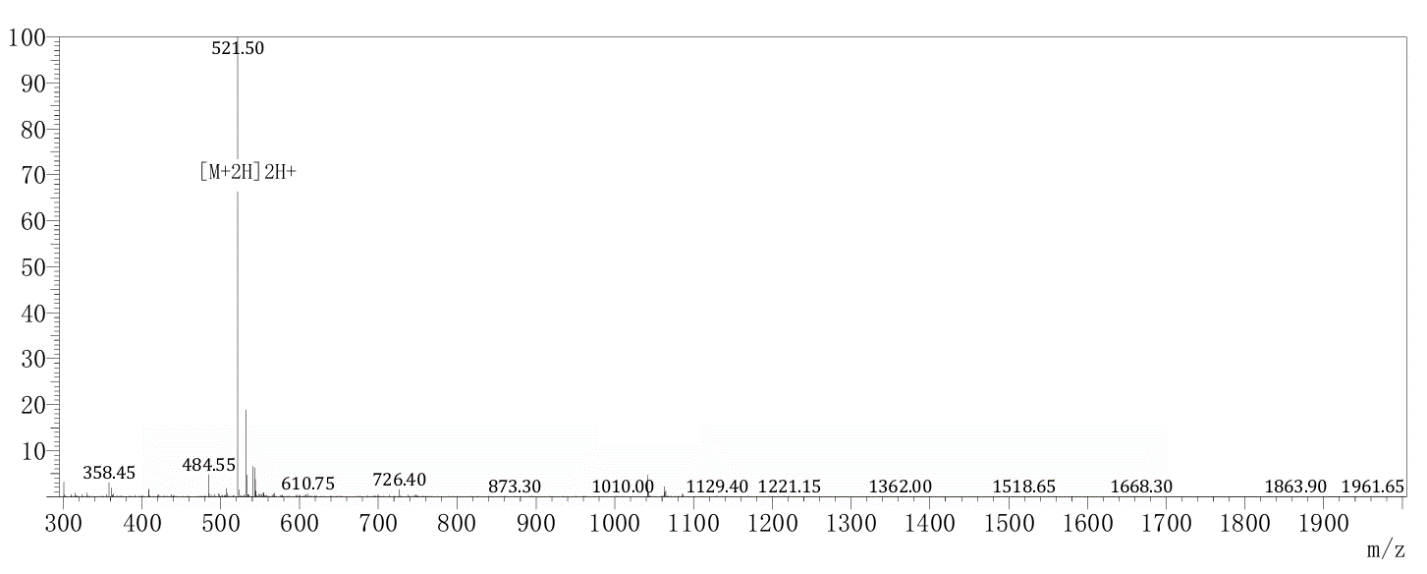

**Figure S4.** LC-MS data for Cyclic peptide 0.

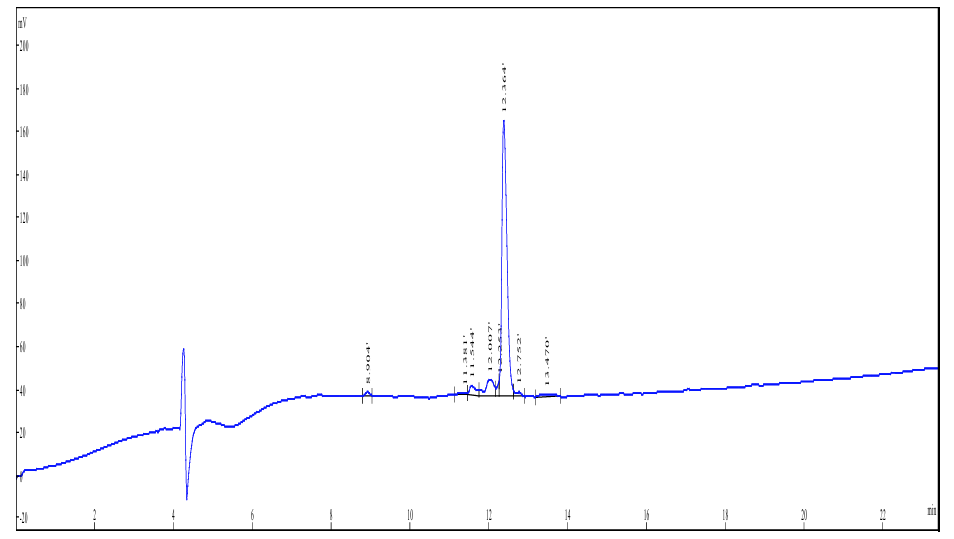

**Figure S5.** HPLC trace for Cyclic peptide 1.

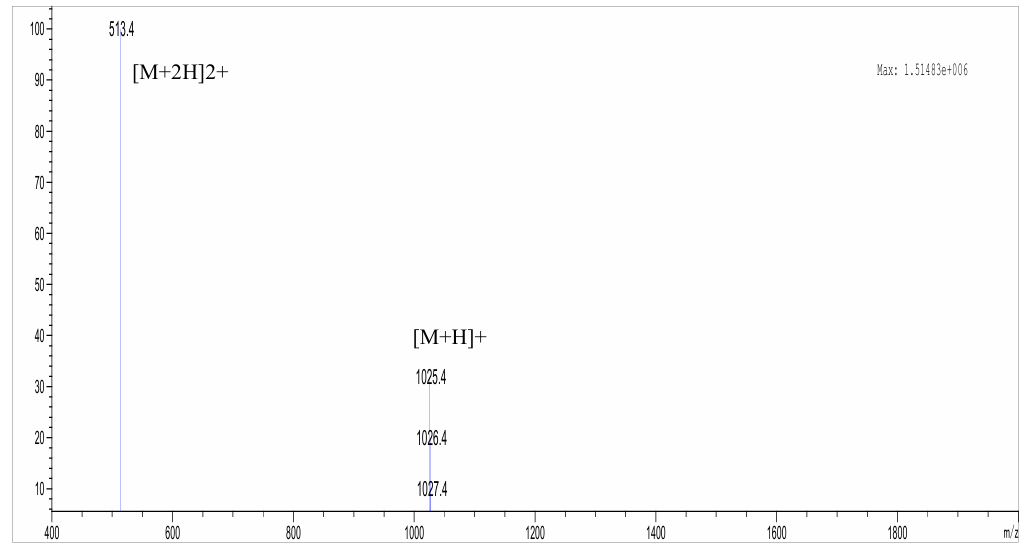

**Figure S6.** LC-MS data for Cyclic peptide 1.

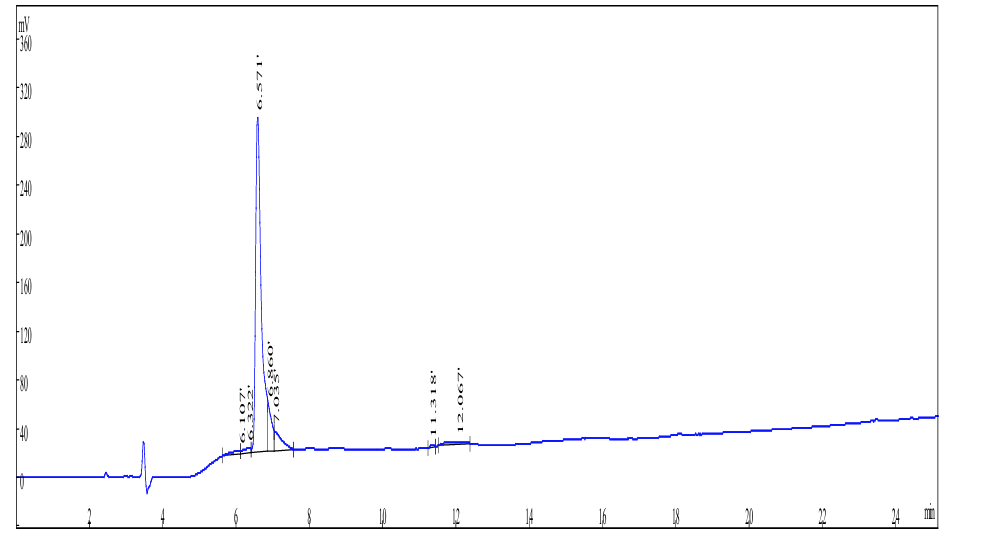

**Figure S7.** HPLC trace for Cyclic peptide 2.

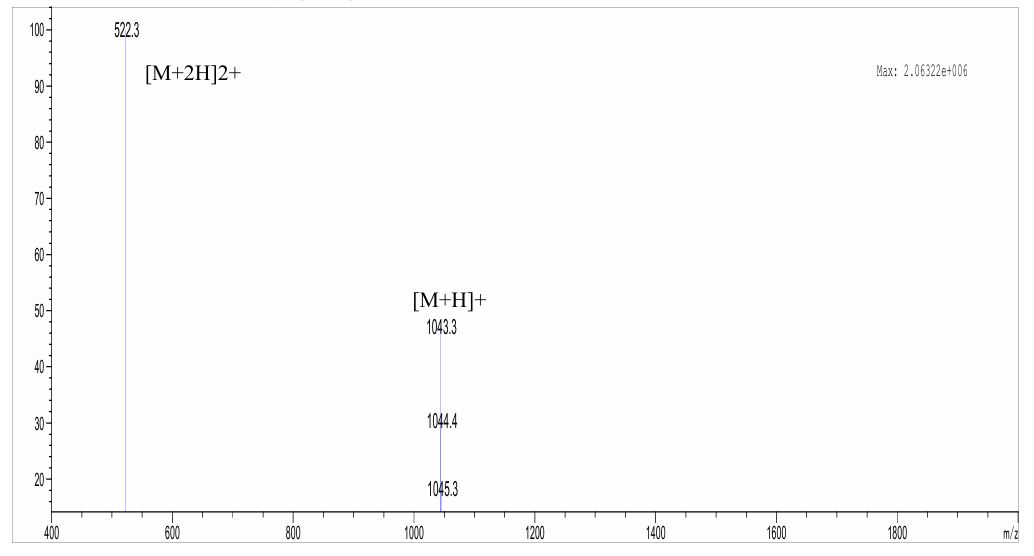

**Figure S8.** LC-MS data for Cyclic peptide 2.

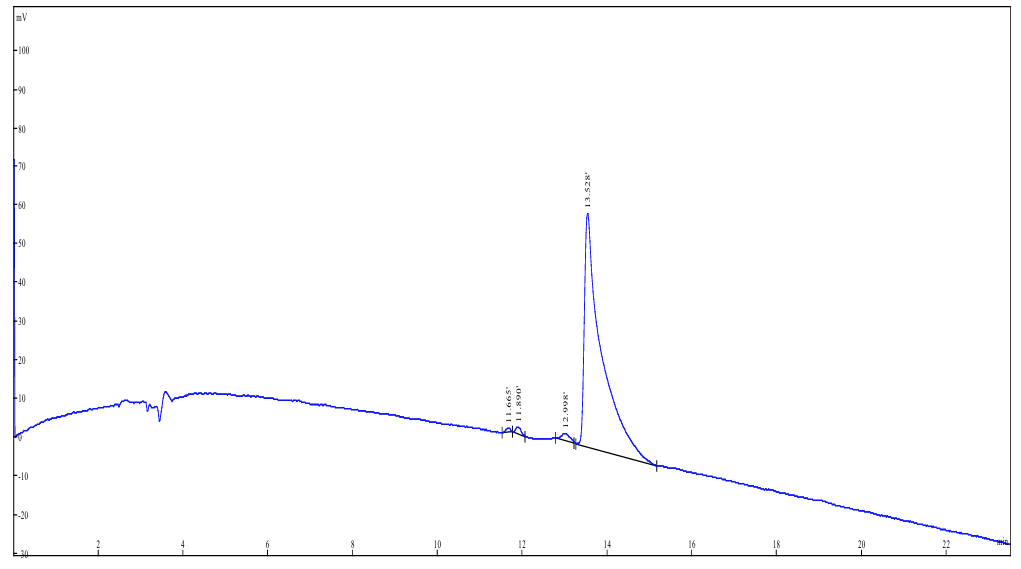

**Figure S9.** HPLC trace for Cyclic peptide 3.

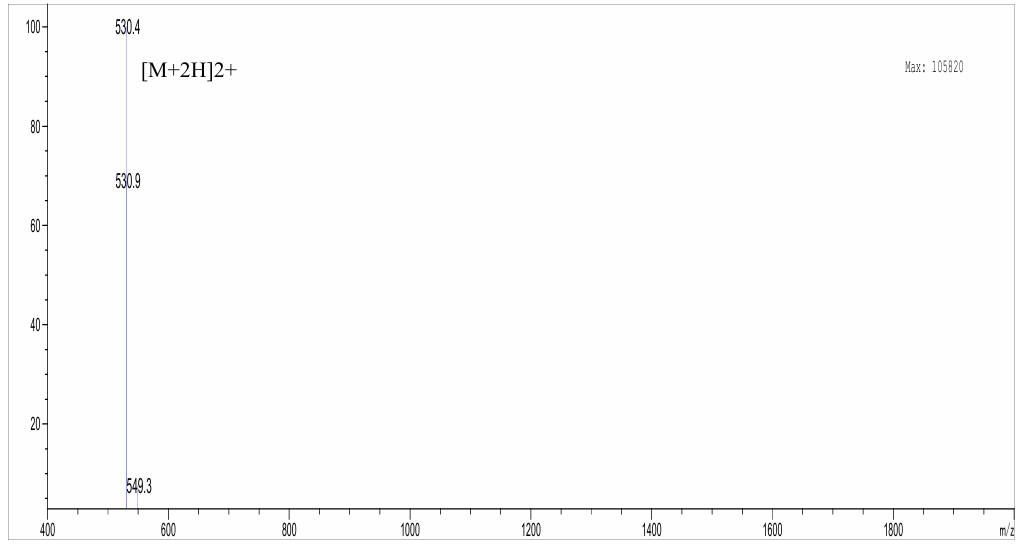

**Figure S10.** LC-MS data for Cyclic peptide 3.

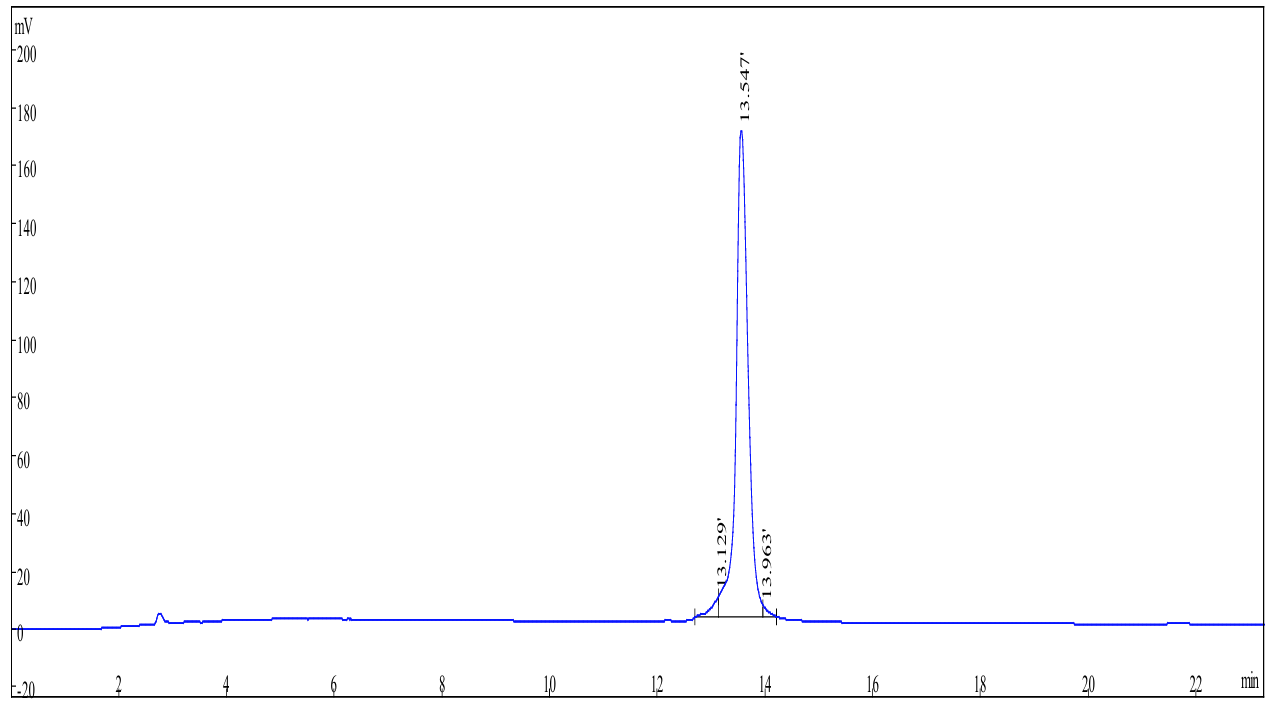

**Figure S11.** HPLC trace for Cyclic peptide 4.

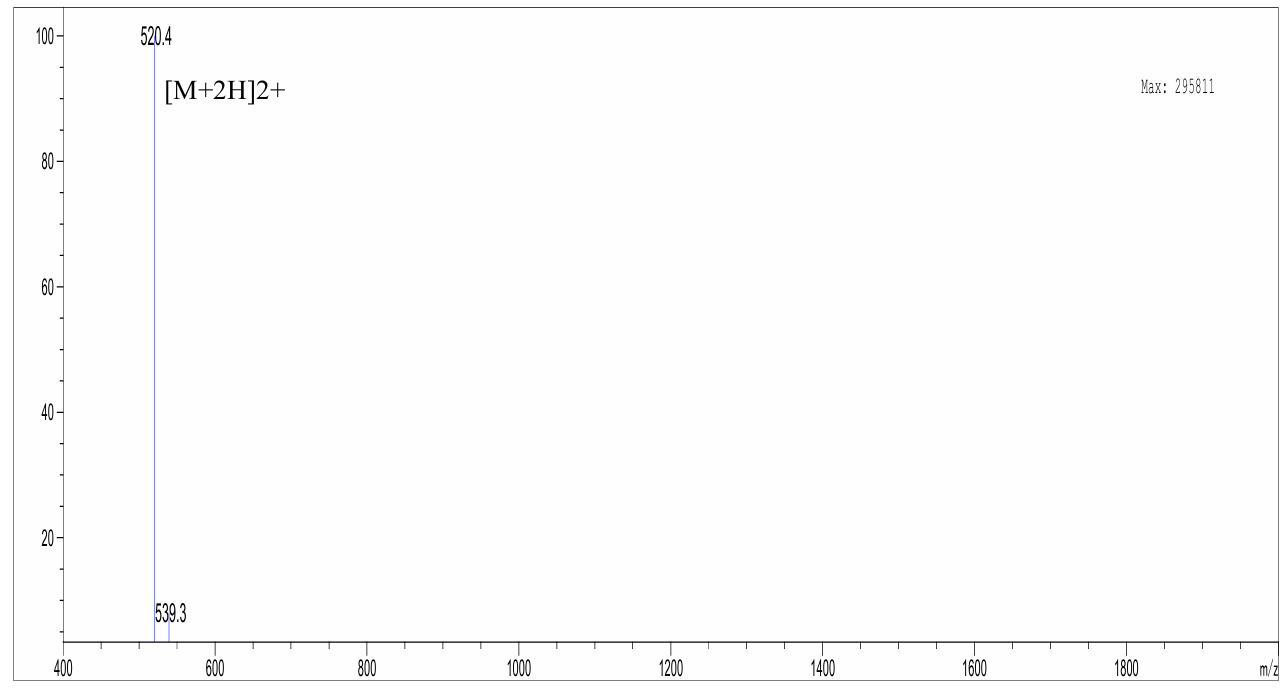

**Figure S12.** LC-MS data for Cyclic peptide 4.

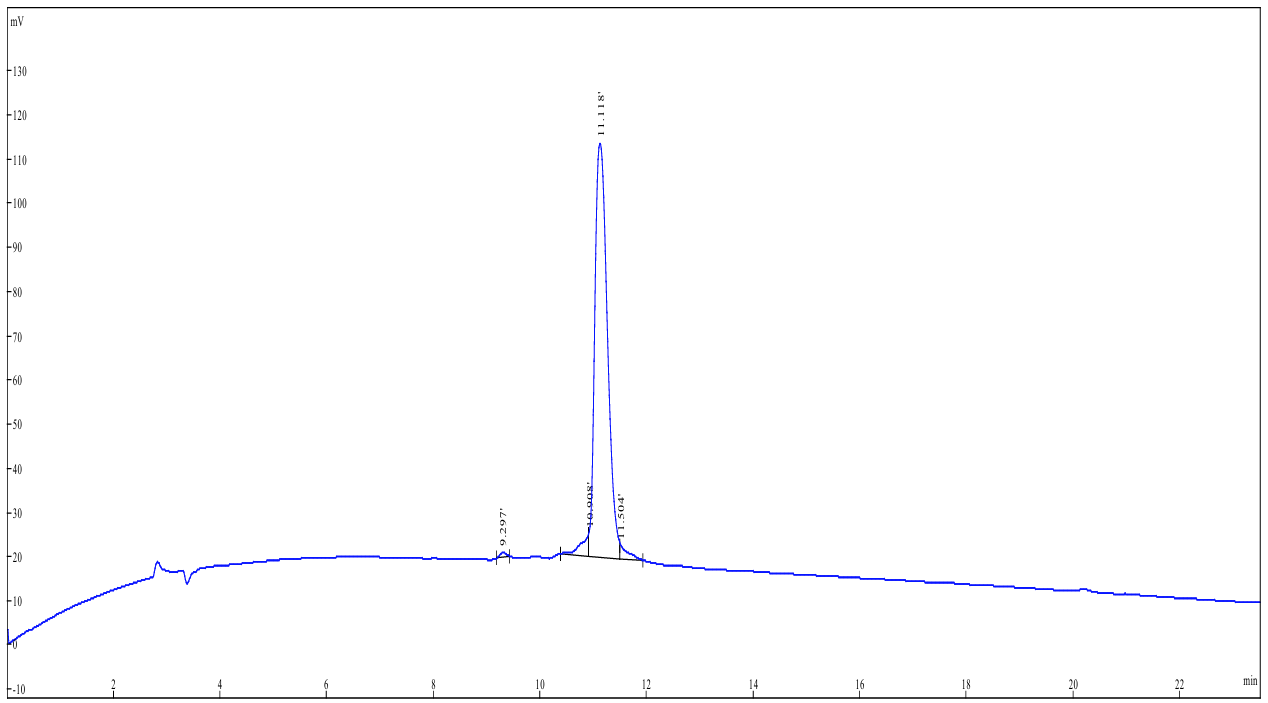

**Figure S13.** HPLC trace for Cyclic peptide 5.

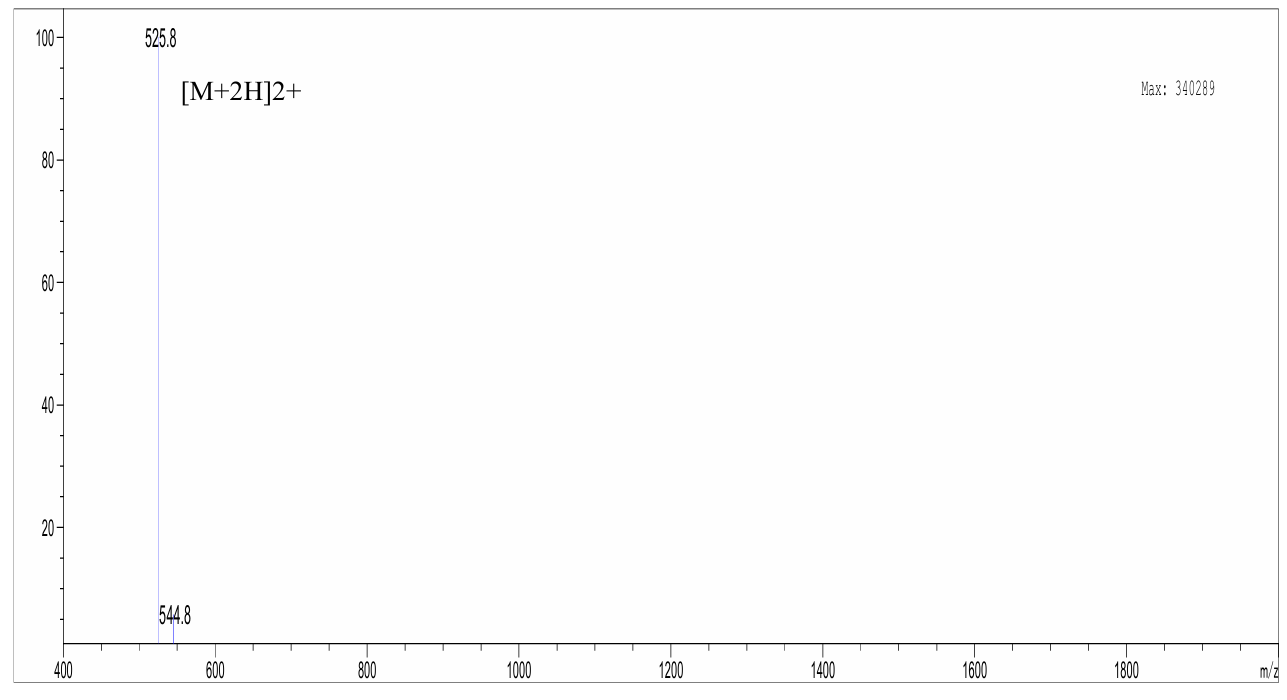

**Figure S14.** LC-MS data for Cyclic peptide 5.

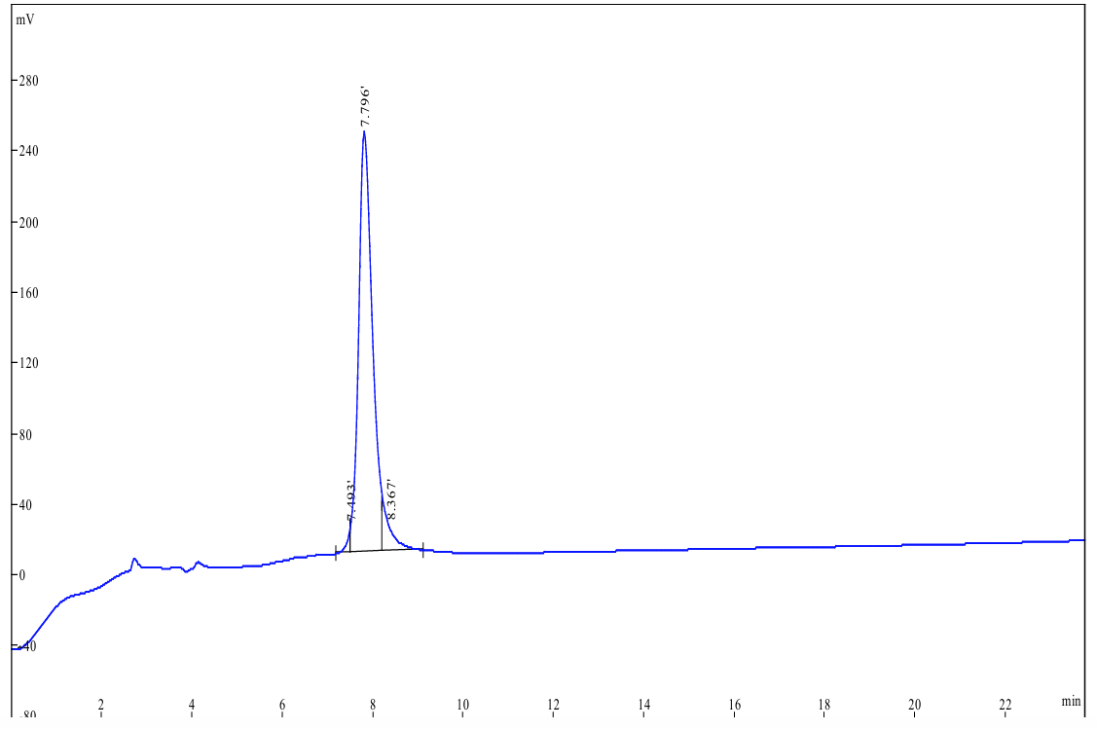

**Figure S15.** HPLC trace for Cyclic peptide 6.

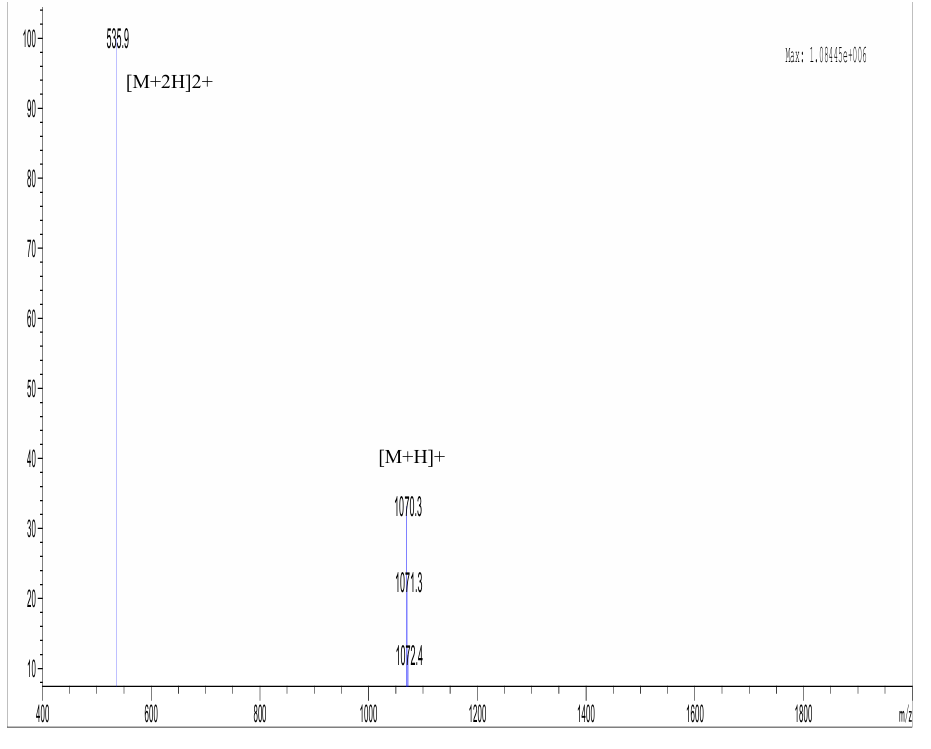

**Figure S16.** LC-MS data for Cyclic peptide 6.

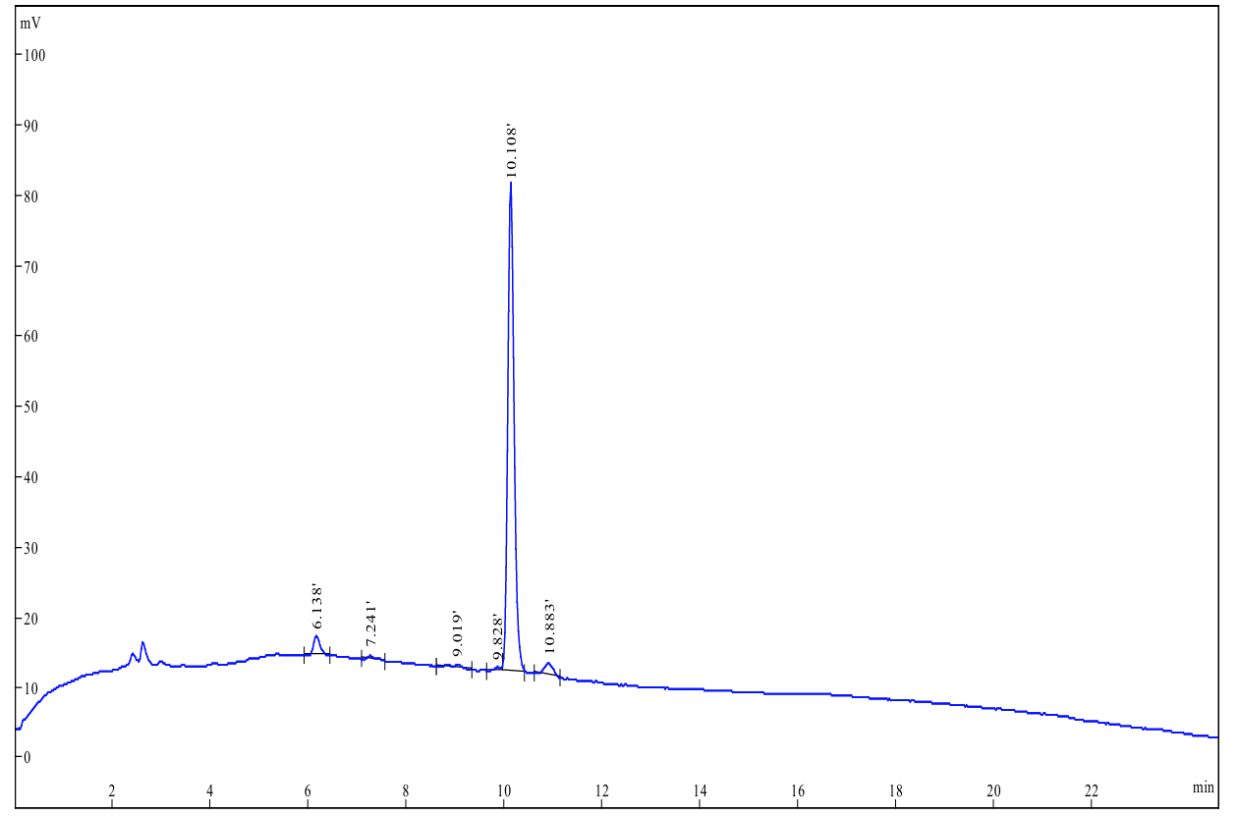

**Figure S17.** HPLC trace for Cyclic peptide 7.

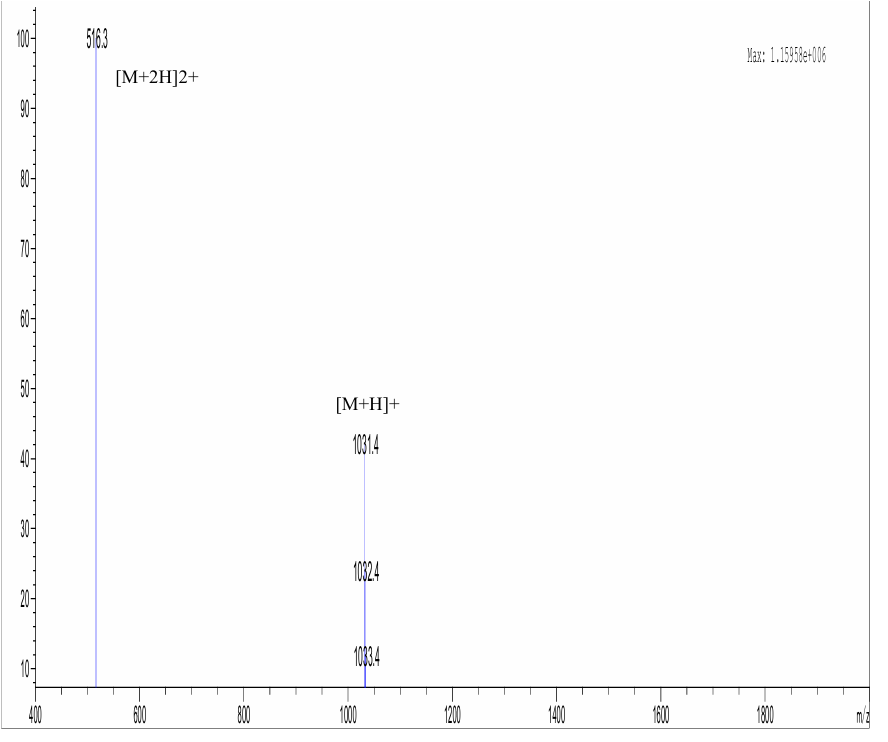

**Figure S18.** LC-MS data for Cyclic peptide 7.

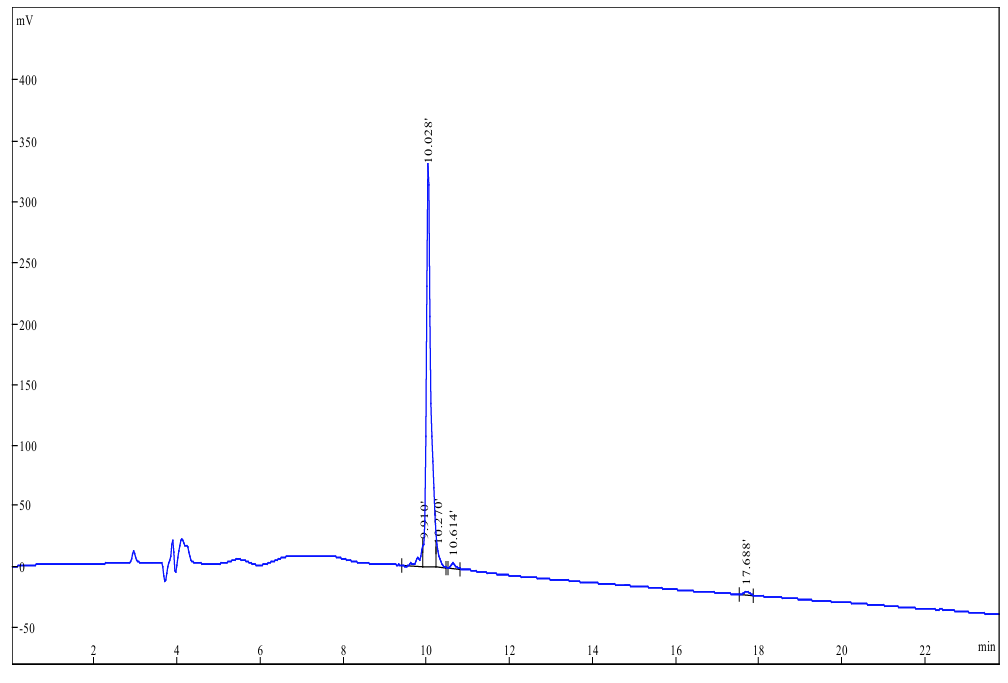

**Figure S19.** HPLC trace for Cyclic peptide 8.

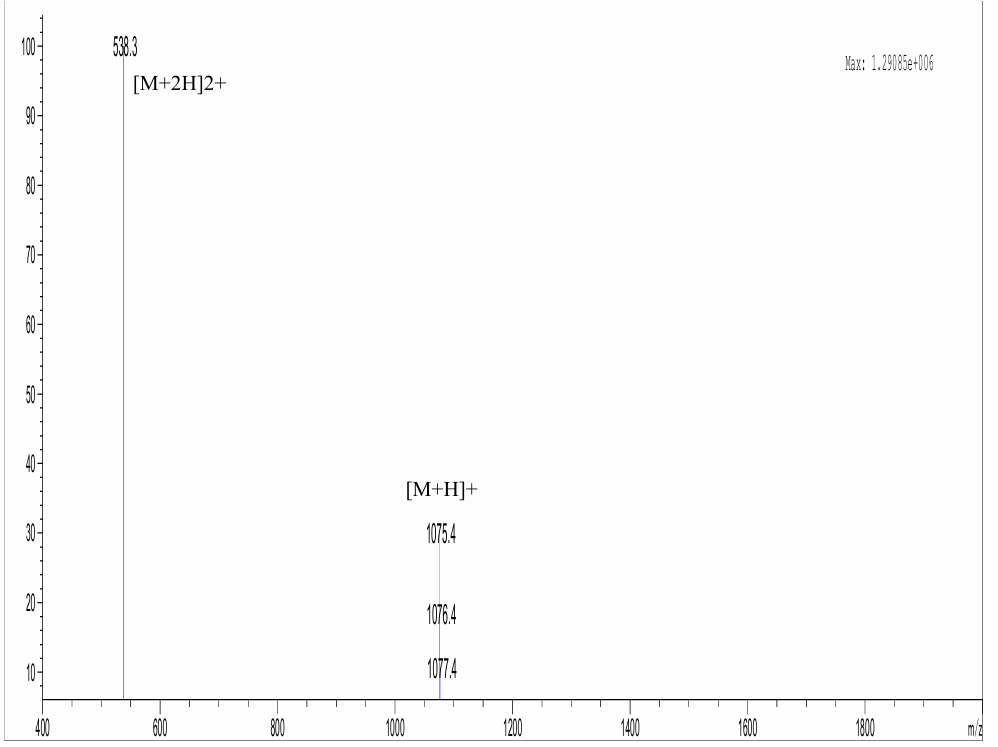

**Figure S20.** LC-MS data for Cyclic peptide 8.

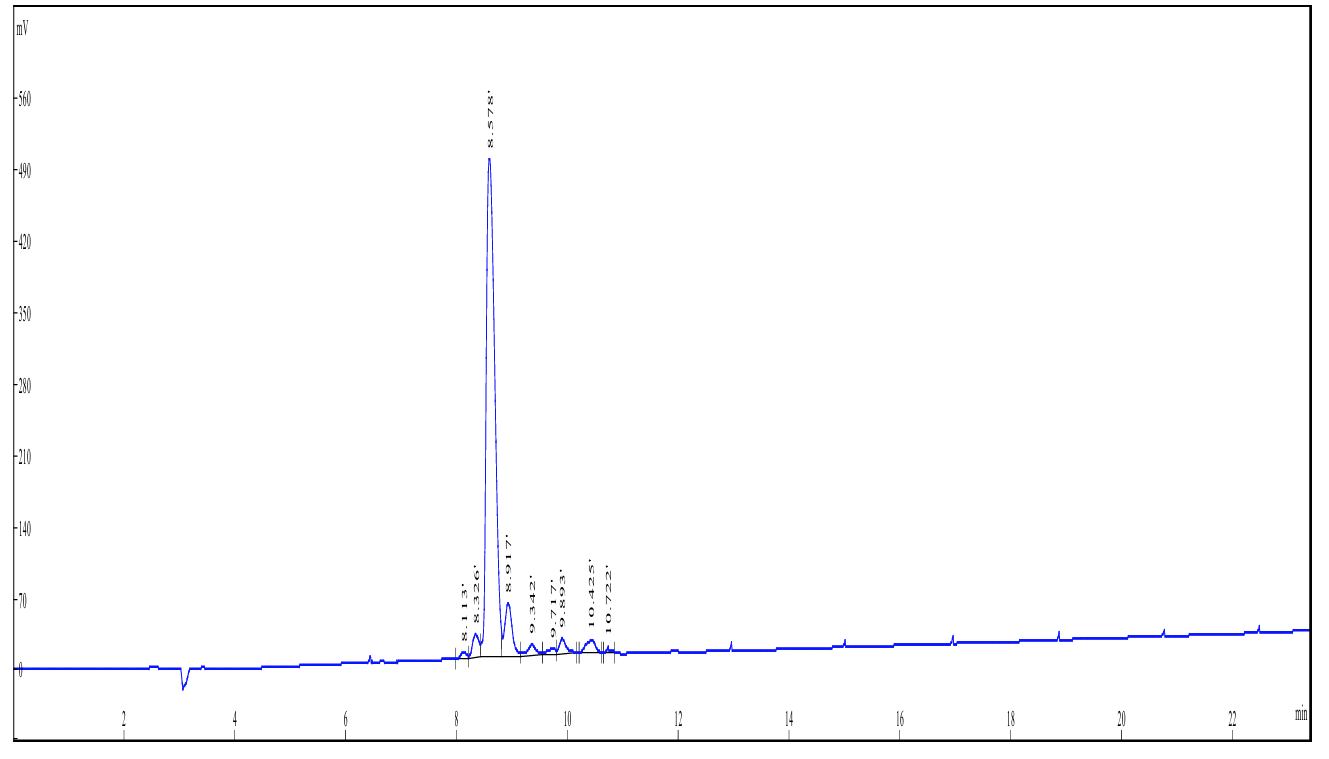

**Figure S21.** HPLC trace for Cyclic peptide 9.

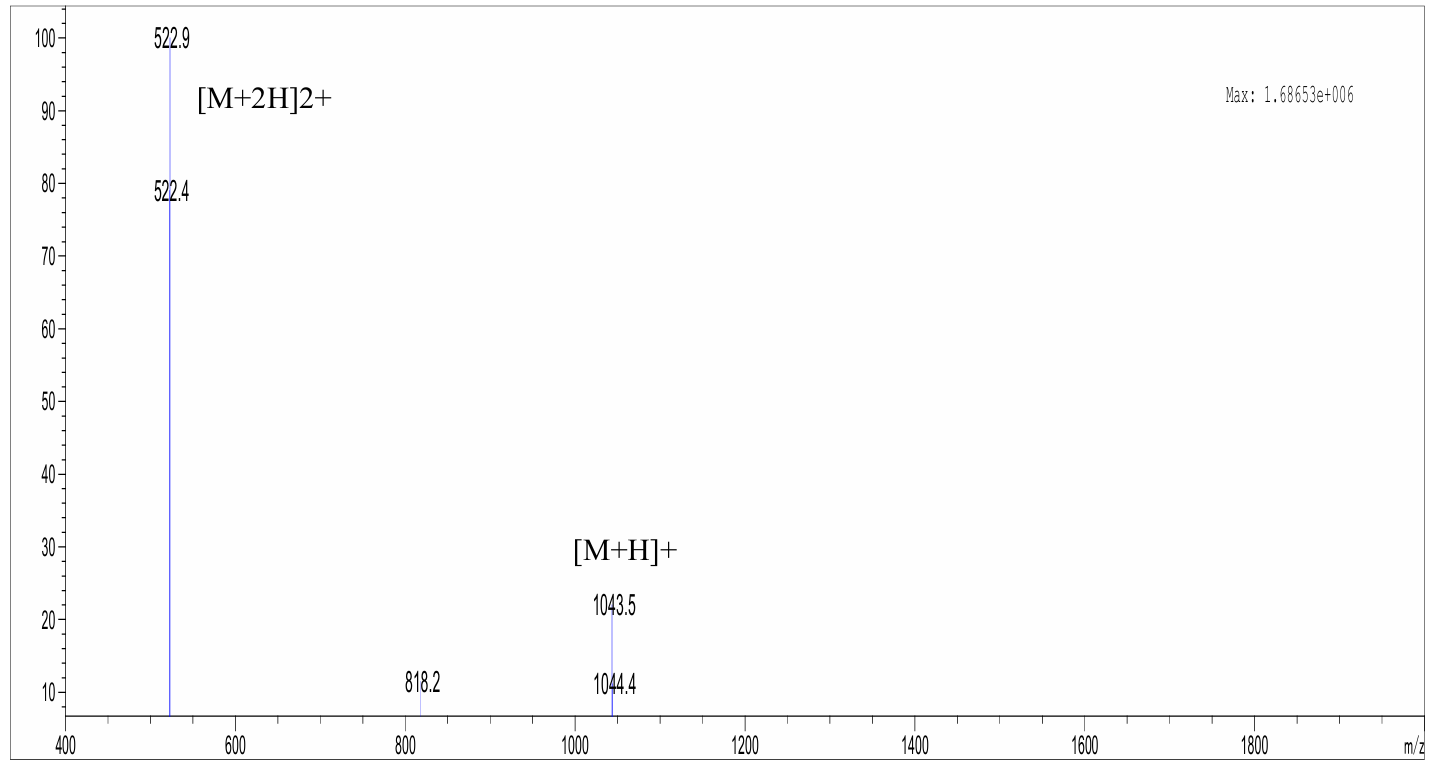

**Figure S22.** LC-MS data for Cyclic peptide 9.

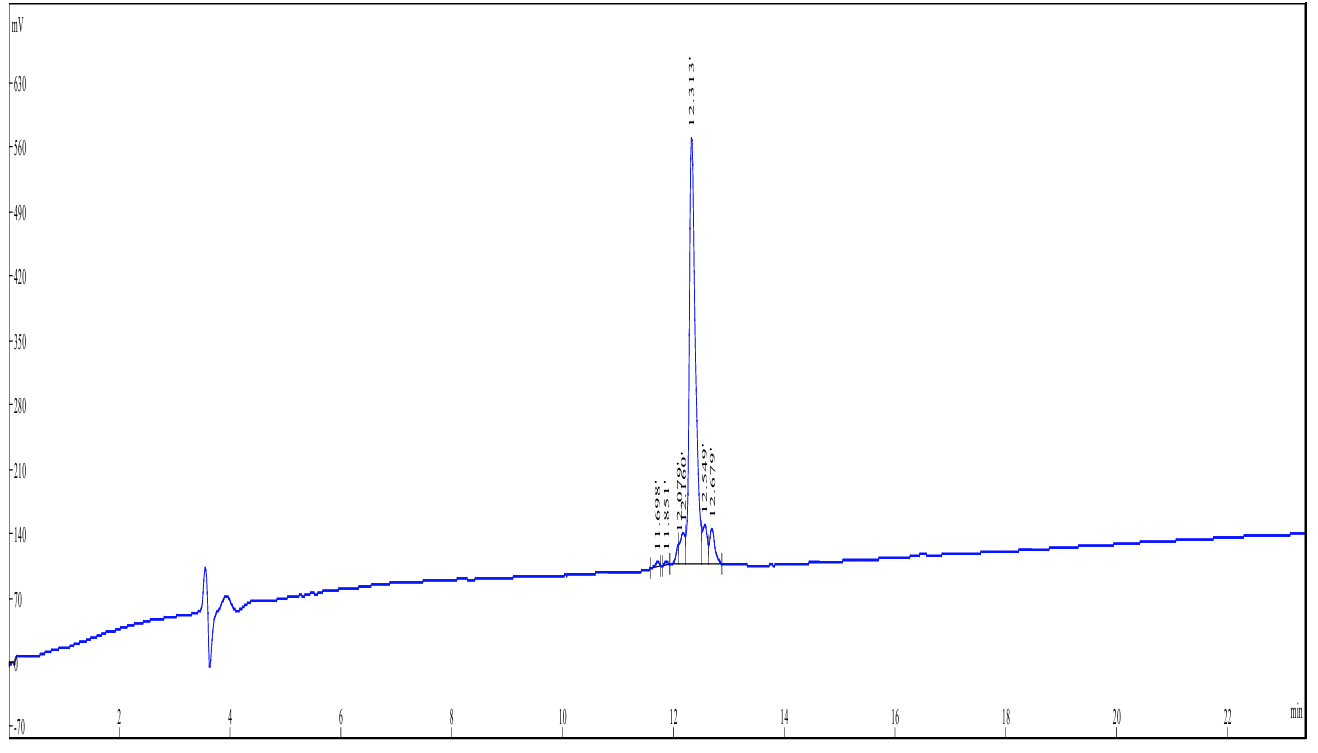

**Figure S23.** HPLC trace for Cyclic peptide 10.

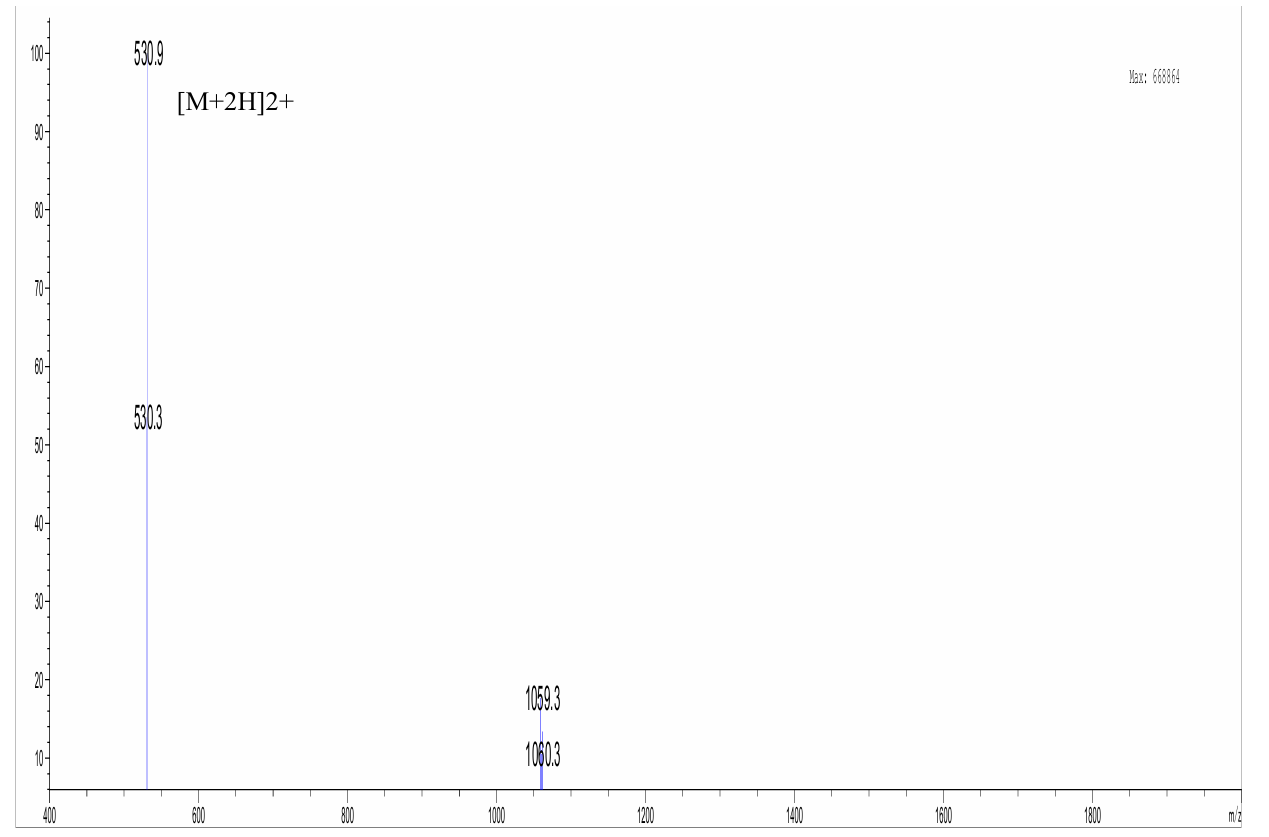

**Figure S24.** LC-MS data for Cyclic peptide 10.

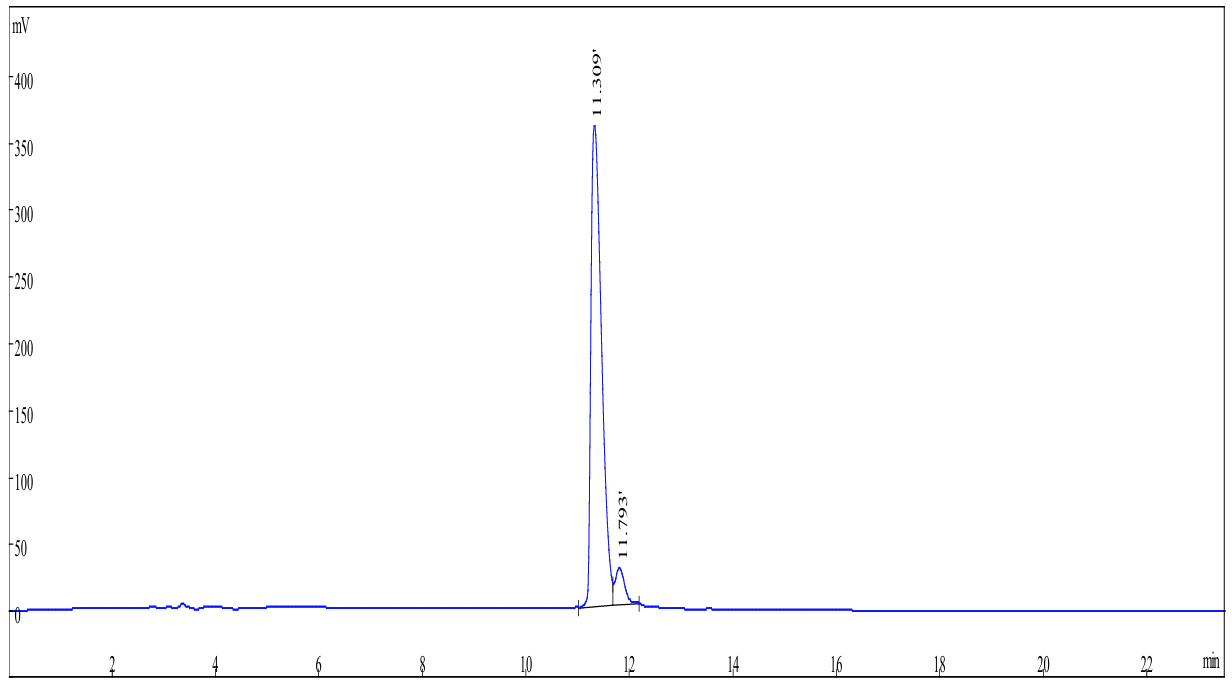

**Figure S25.** HPLC trace for Cyclic peptide 11.

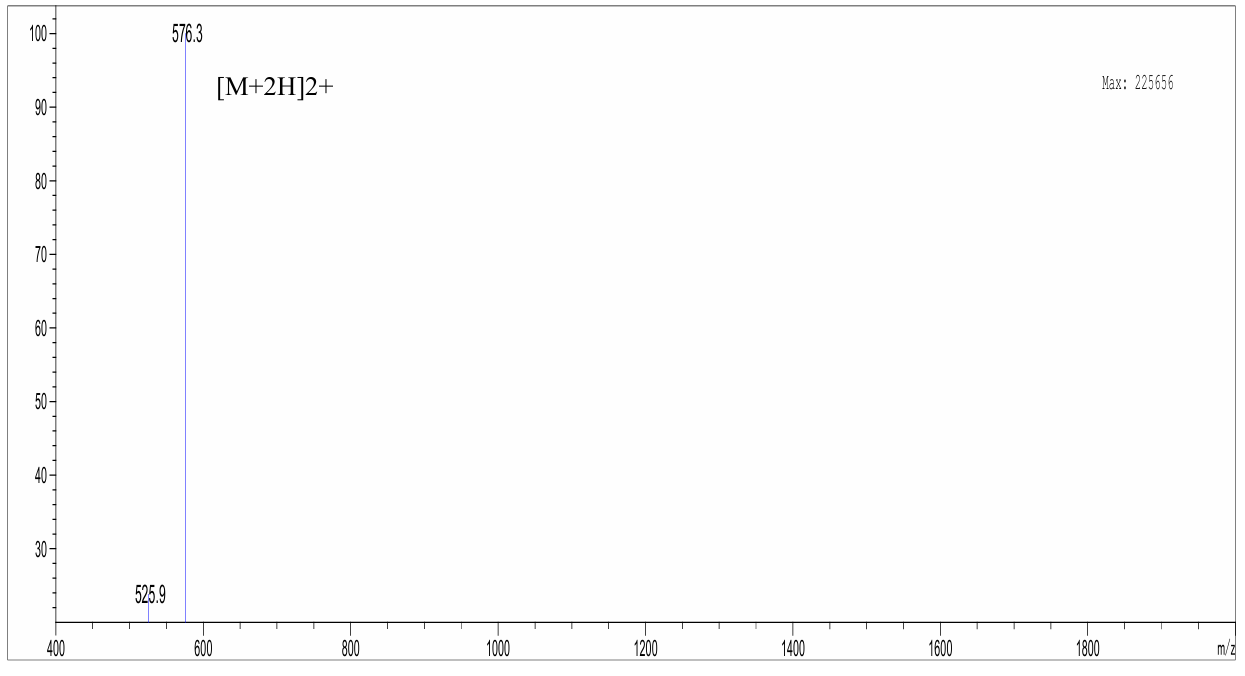

**Figure S26.** LC-MS data for Cyclic peptide 11.

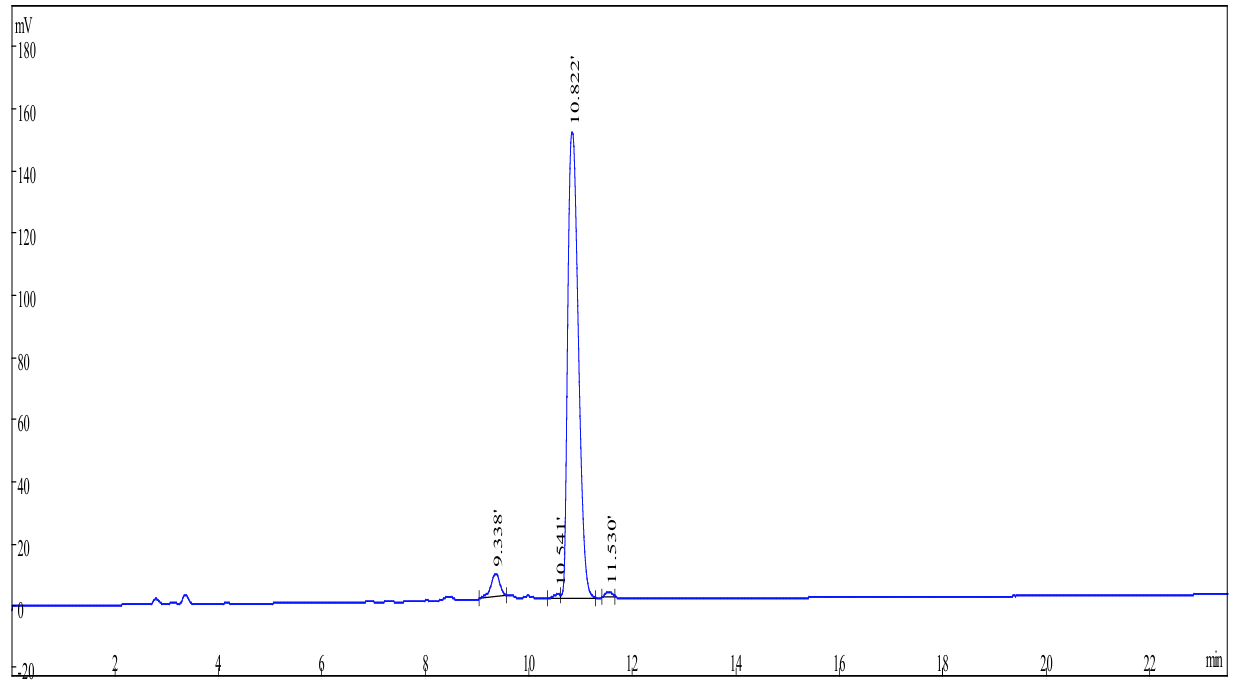

**Figure S27.** HPLC trace for Cyclic peptide 12.

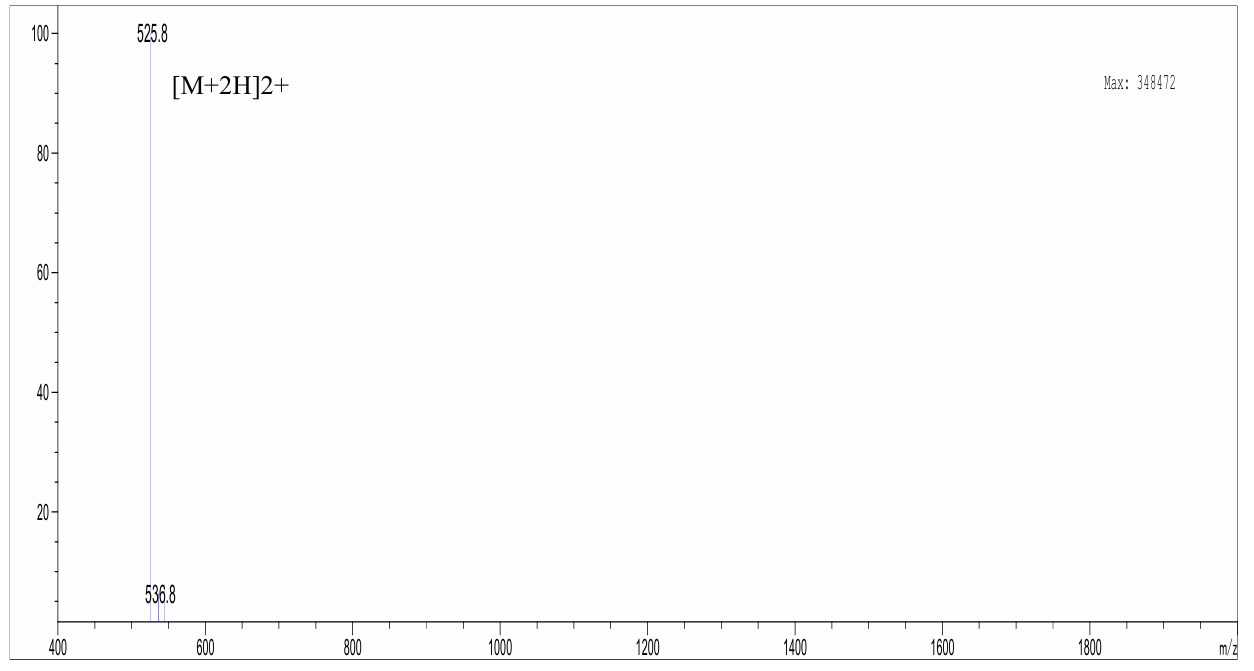

**Figure S28.** LC-MS data for Cyclic peptide 12.

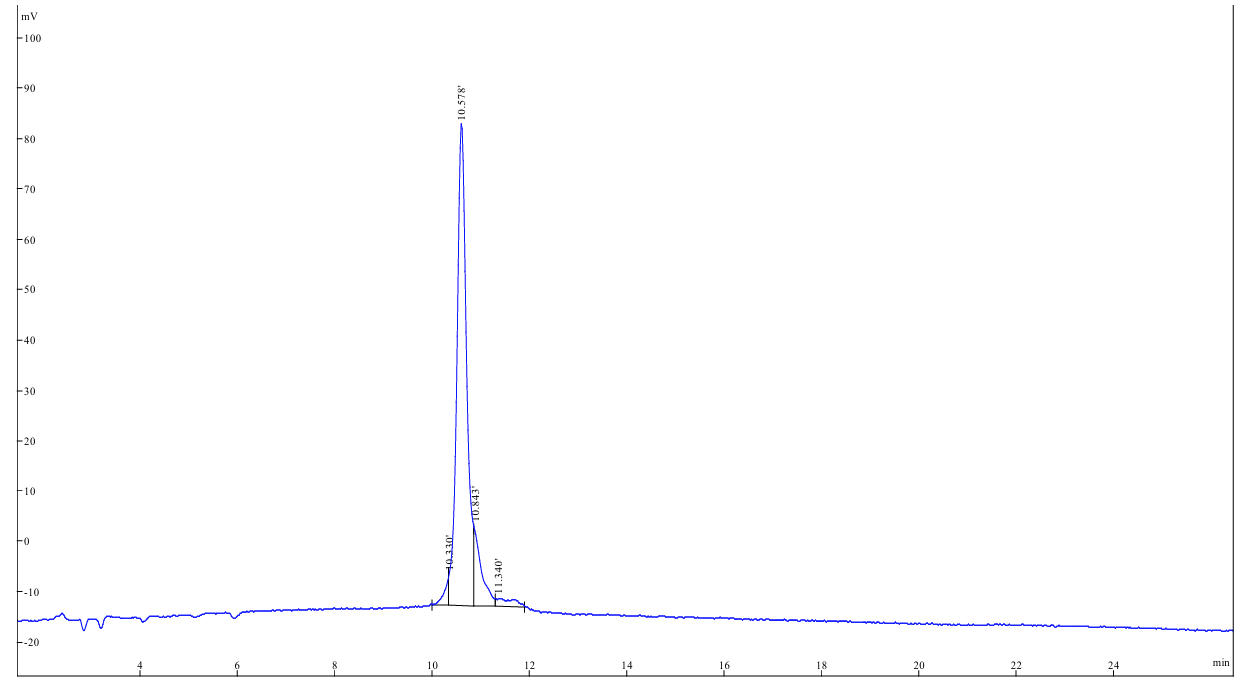

**Figure S29.** HPLC trace for Cyclic peptide 13.

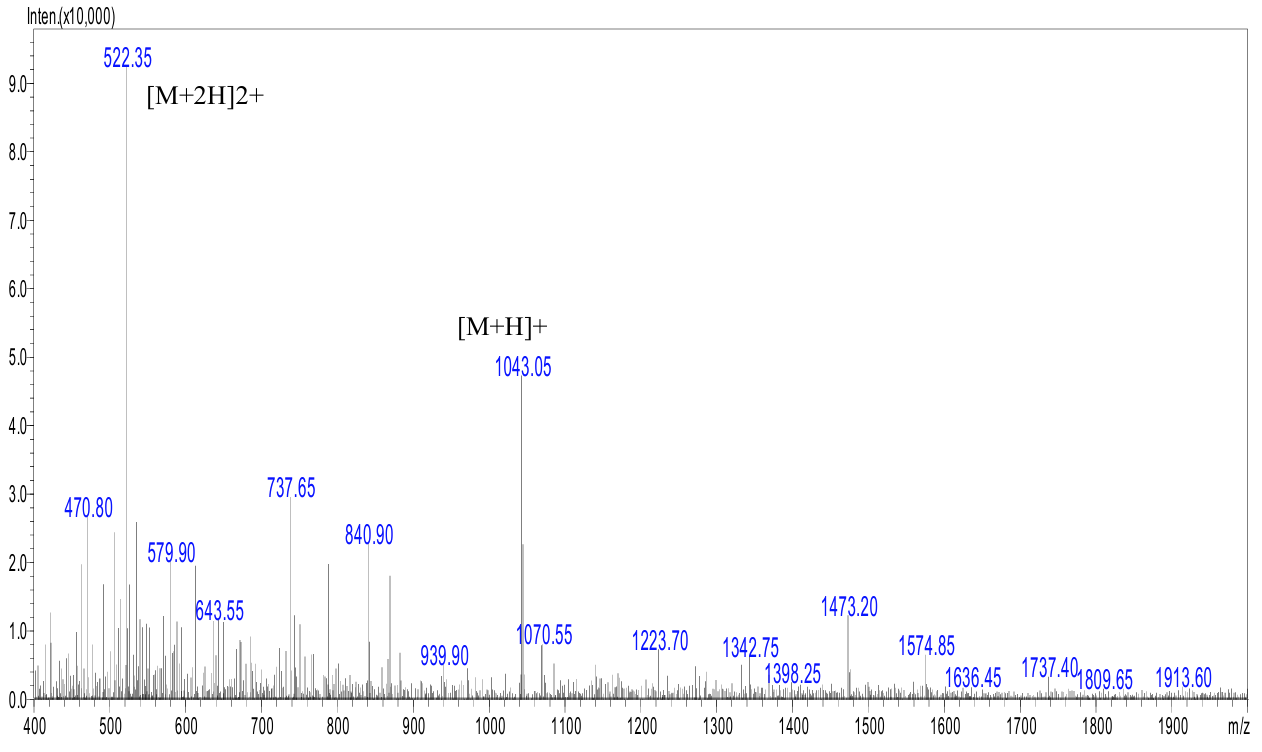

**Figure S30.** LC-MS data for Cyclic peptide 13.

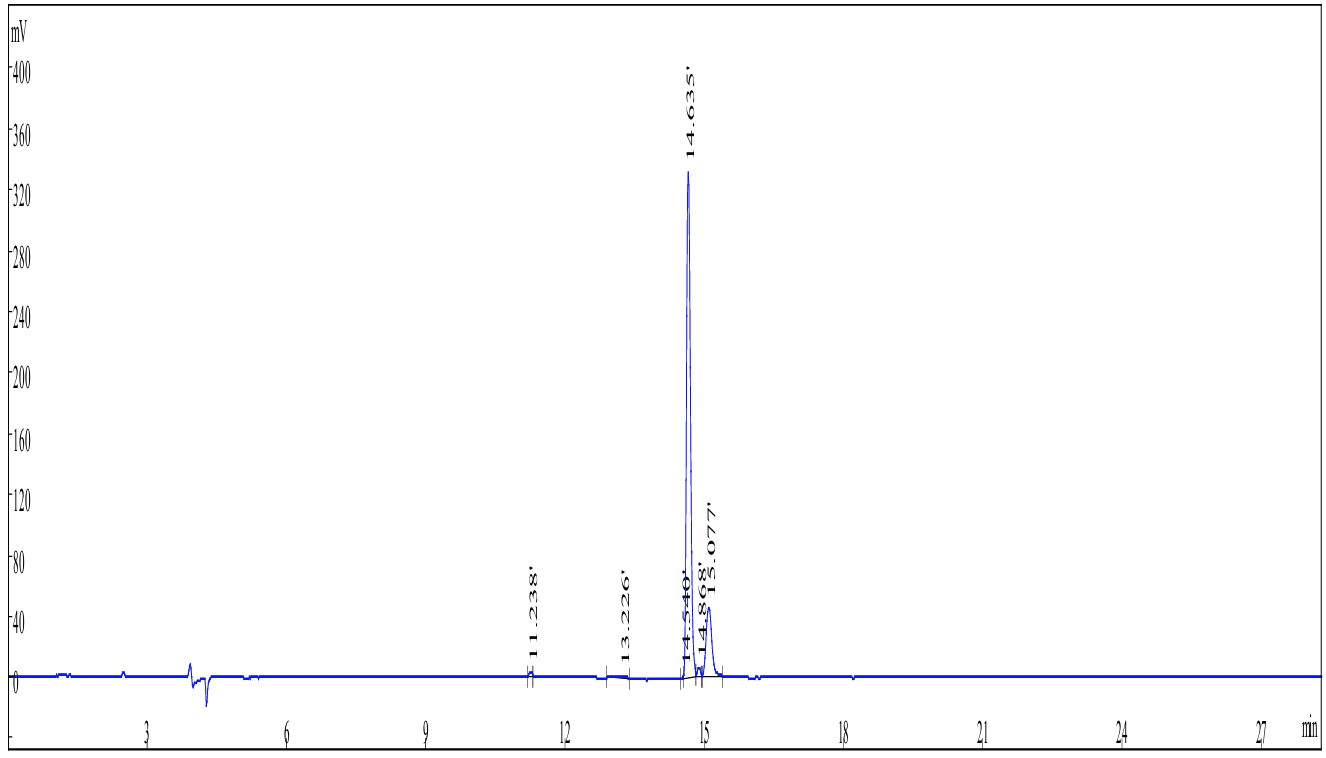

**Figure S31.** HPLC trace for Cyclic peptide 14.

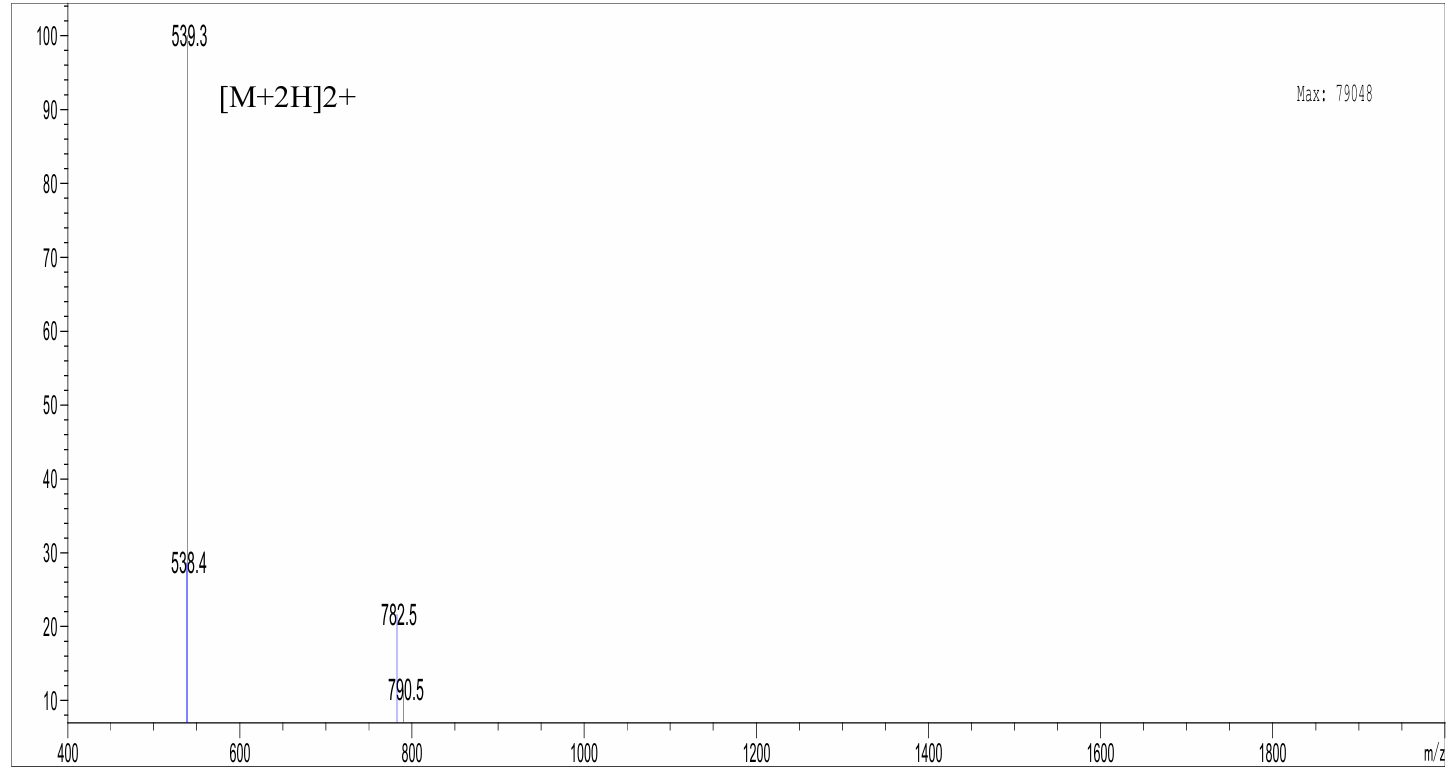

**Figure S32.** LC-MS data for Cyclic peptide 14.

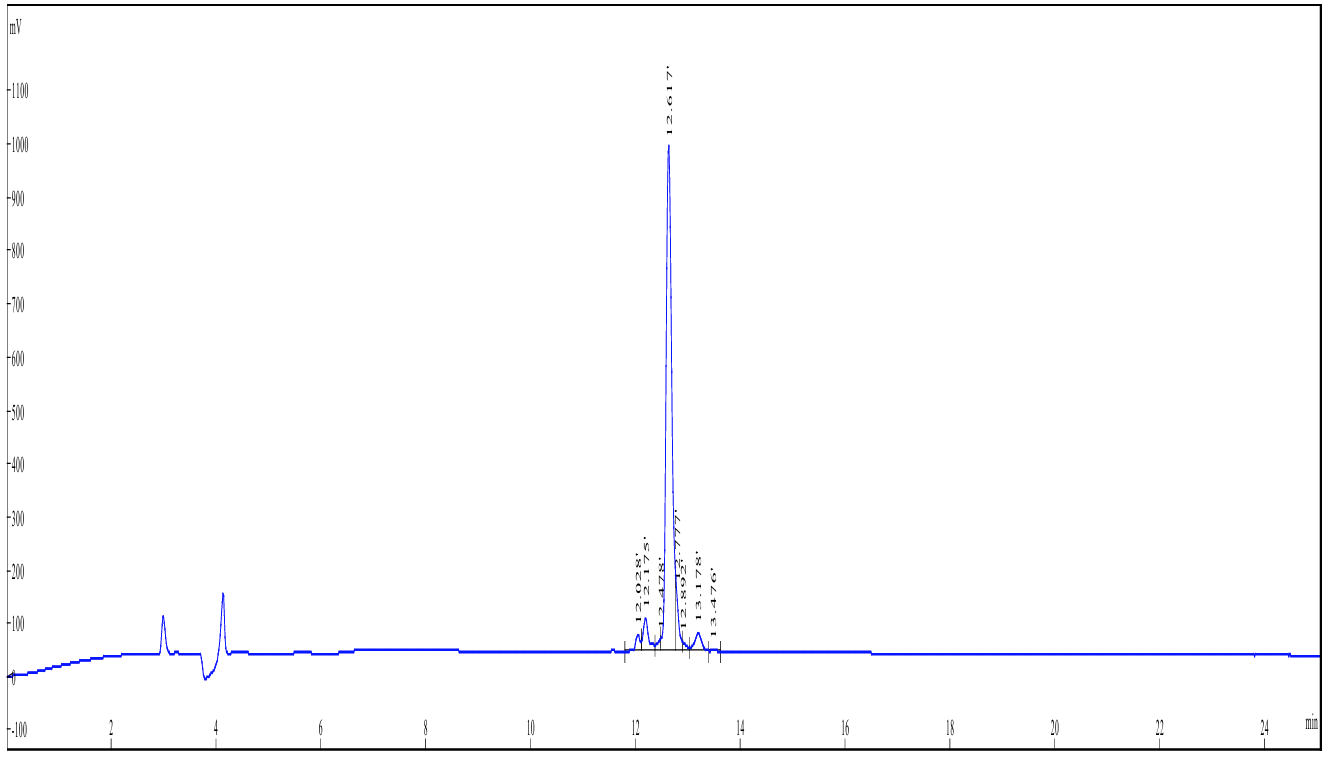

**Figure S33.** HPLC trace for Cyclic peptide 15.

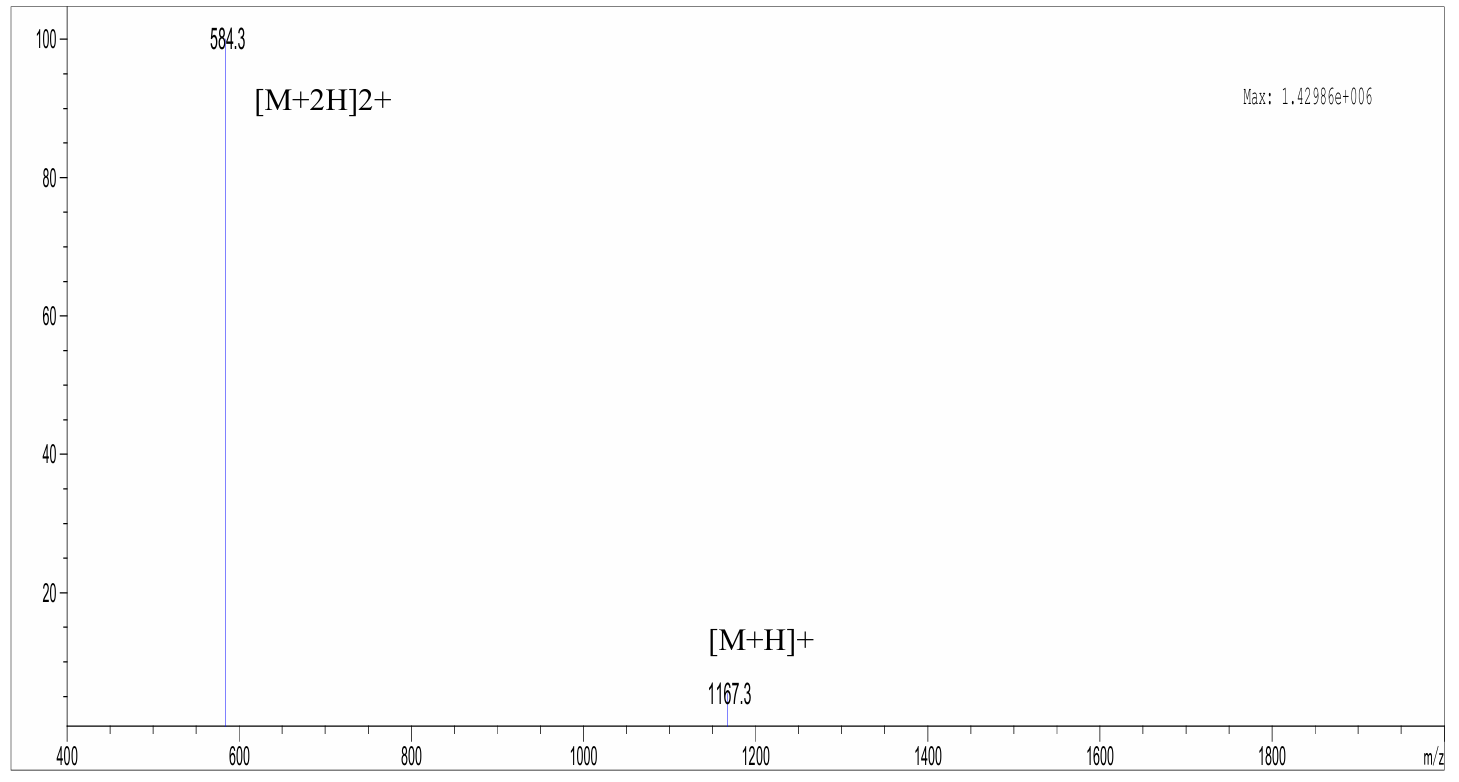

**Figure S34.** LC-MS data for Cyclic peptide 15.

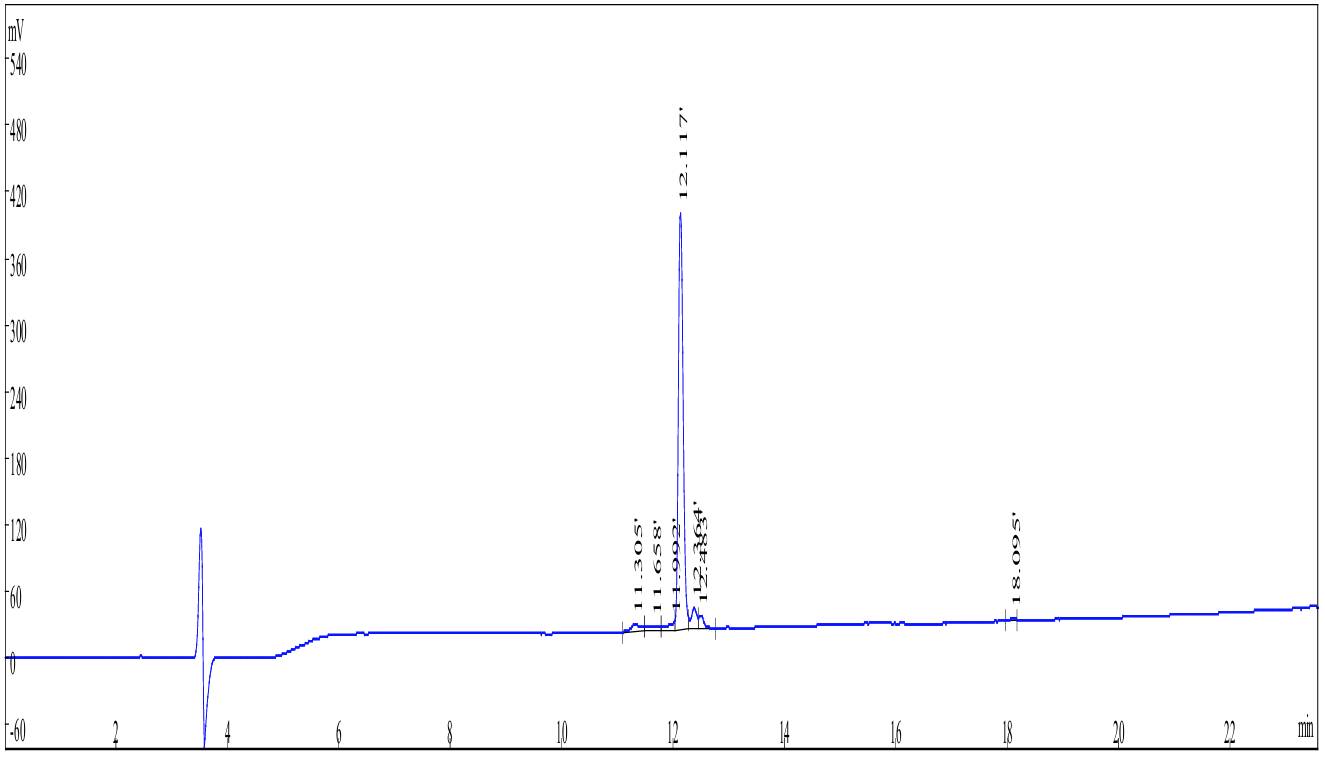

**Figure S35.** HPLC trace for Cyclic peptide 16.

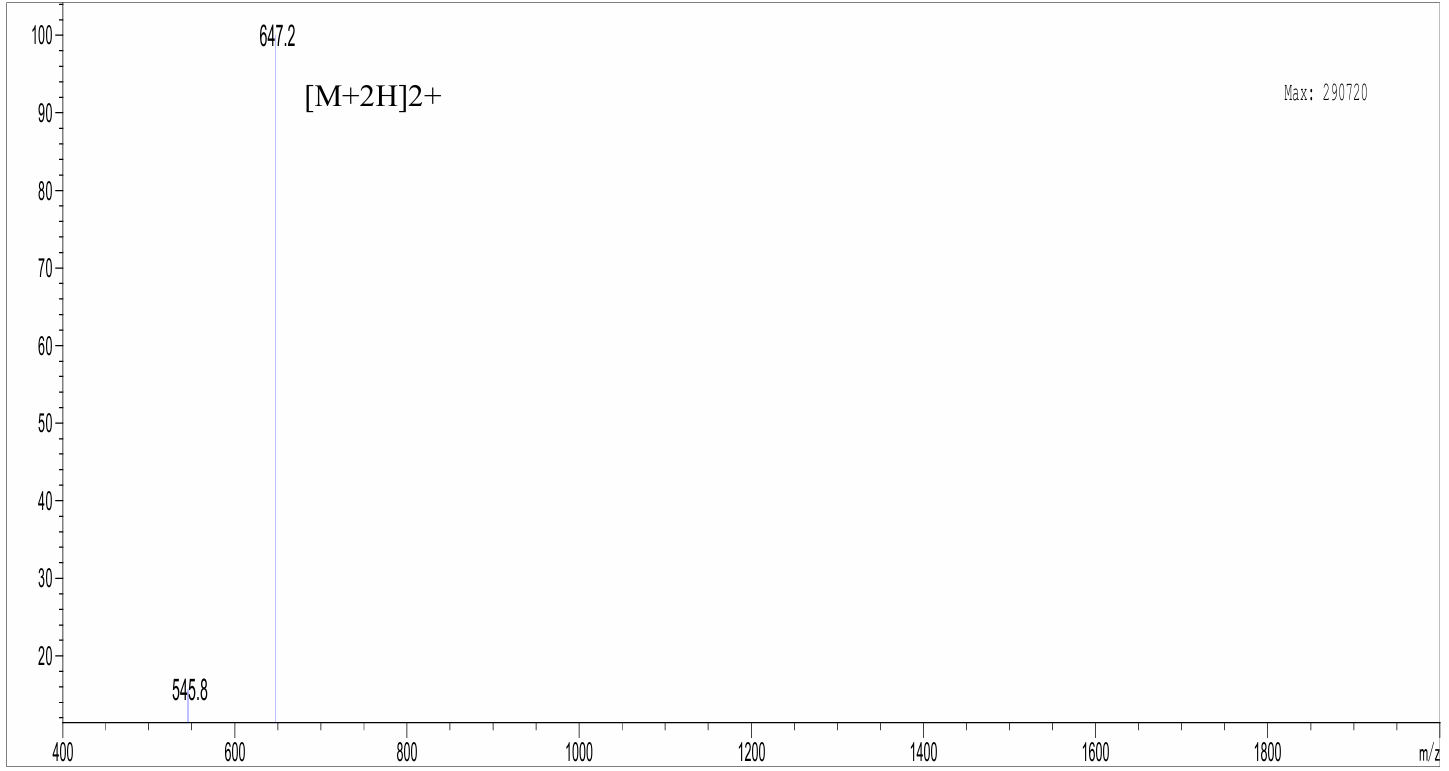

**Figure S36.** LC-MS data for Cyclic peptide 16.

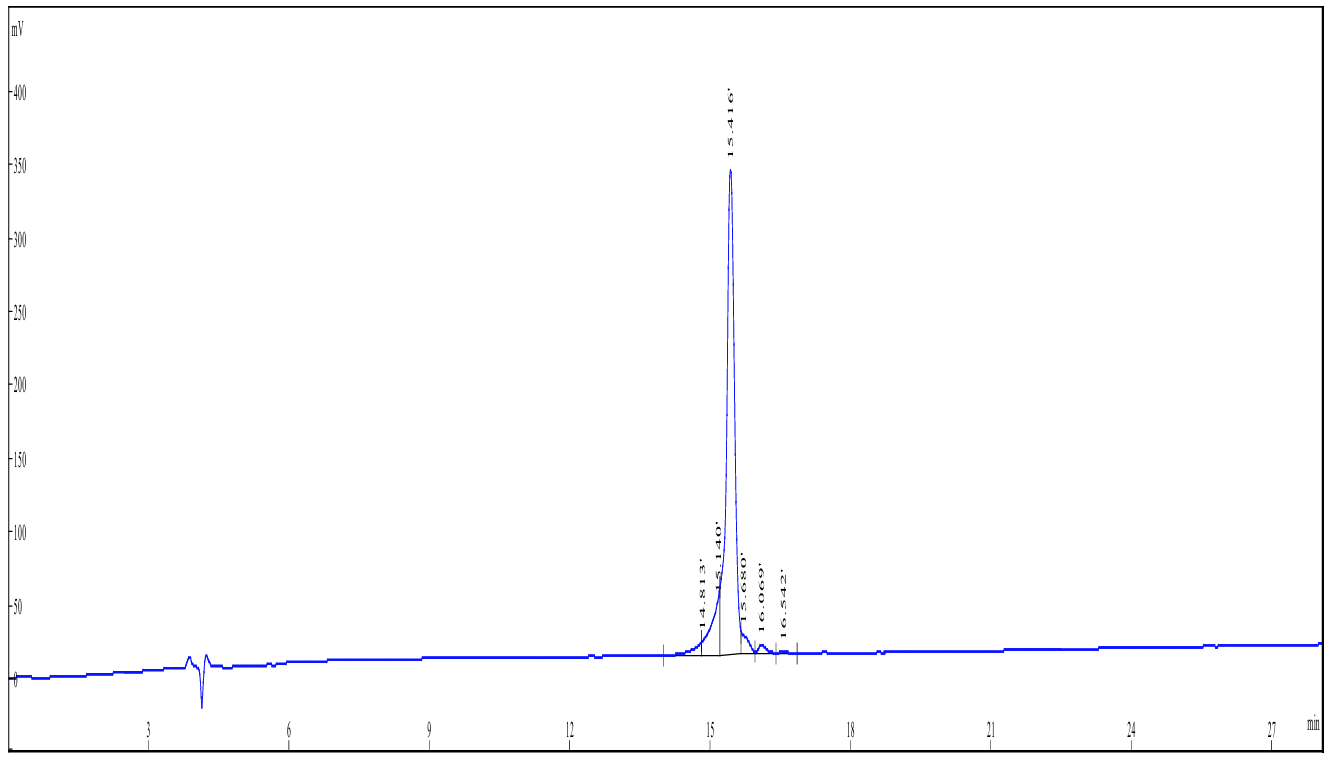

**Figure S37.** HPLC trace for Cyclic peptide 17.

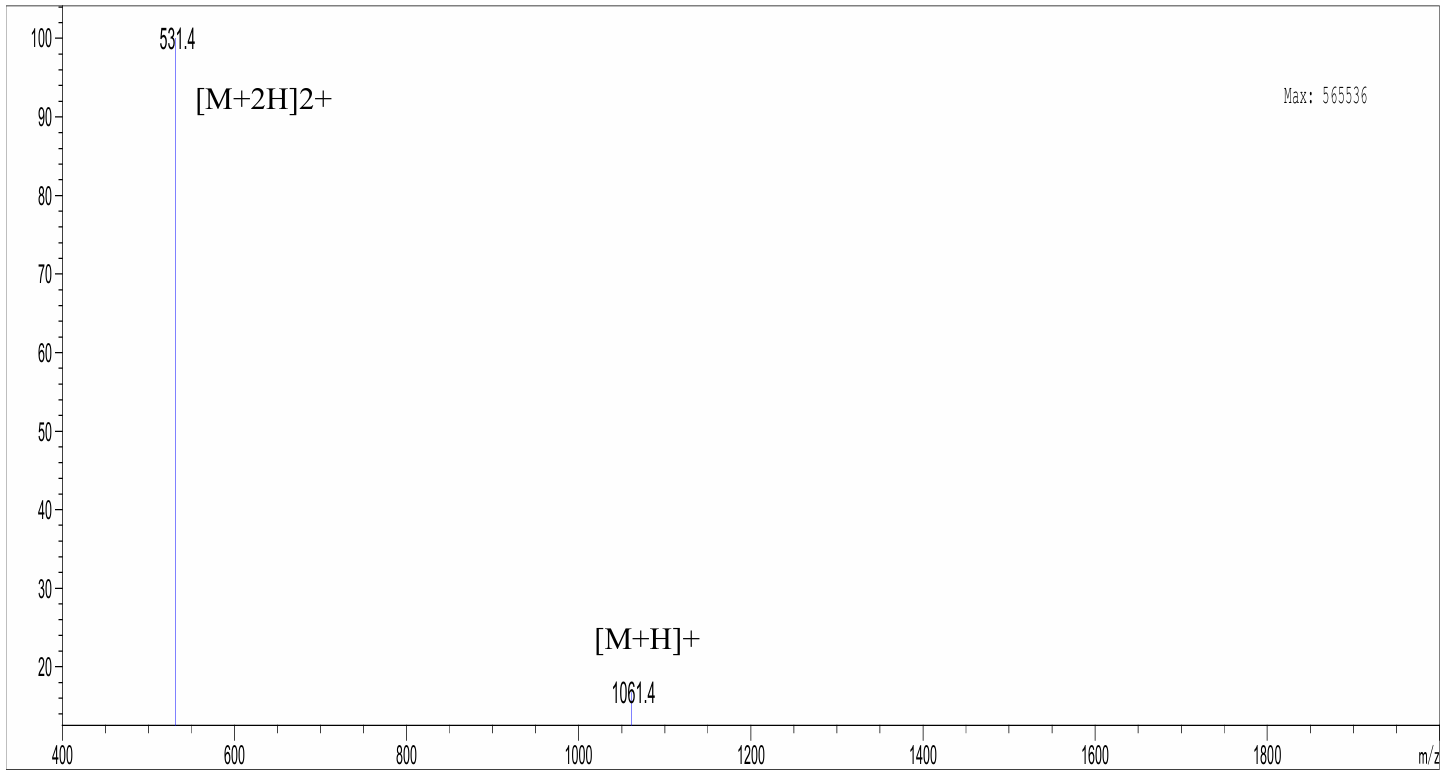

**Figure S38.** LC-MS data for Cyclic peptide 17.

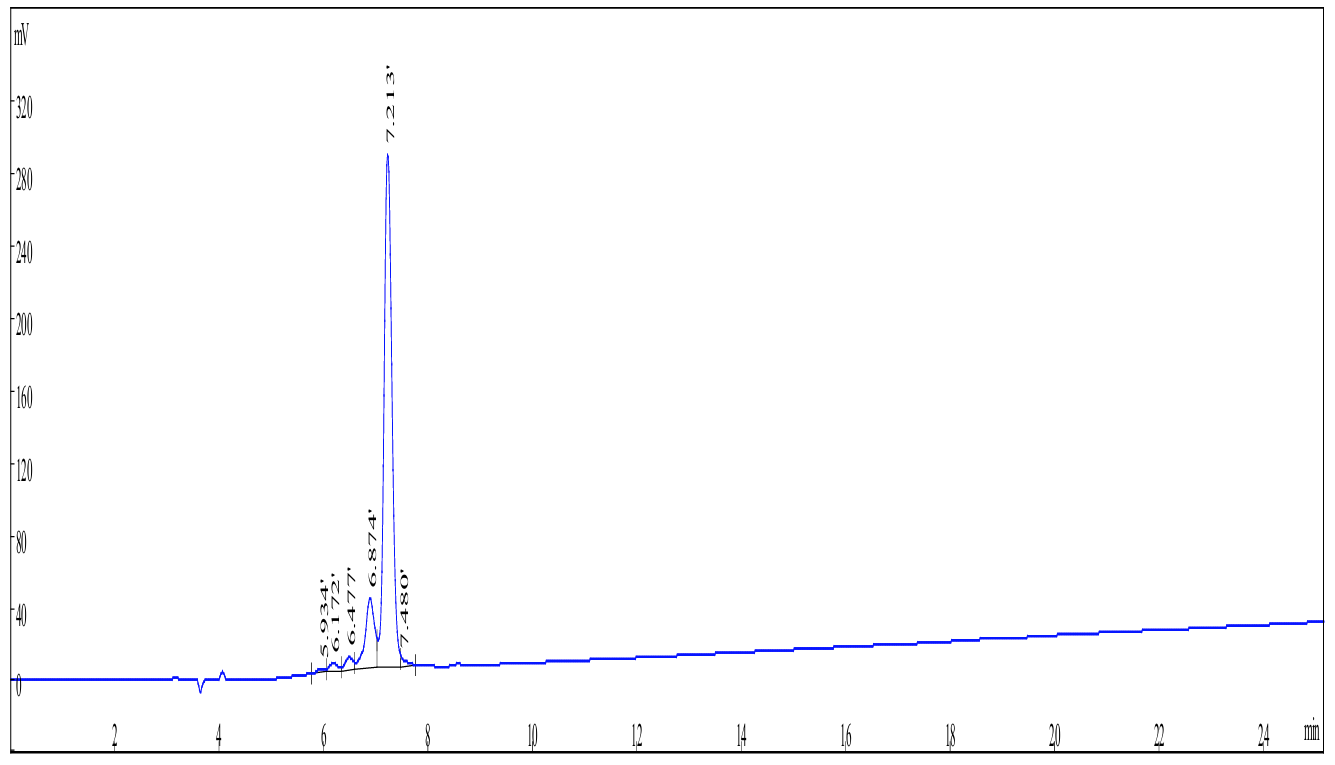

**Figure S39.** HPLC trace for Cyclic peptide 18.

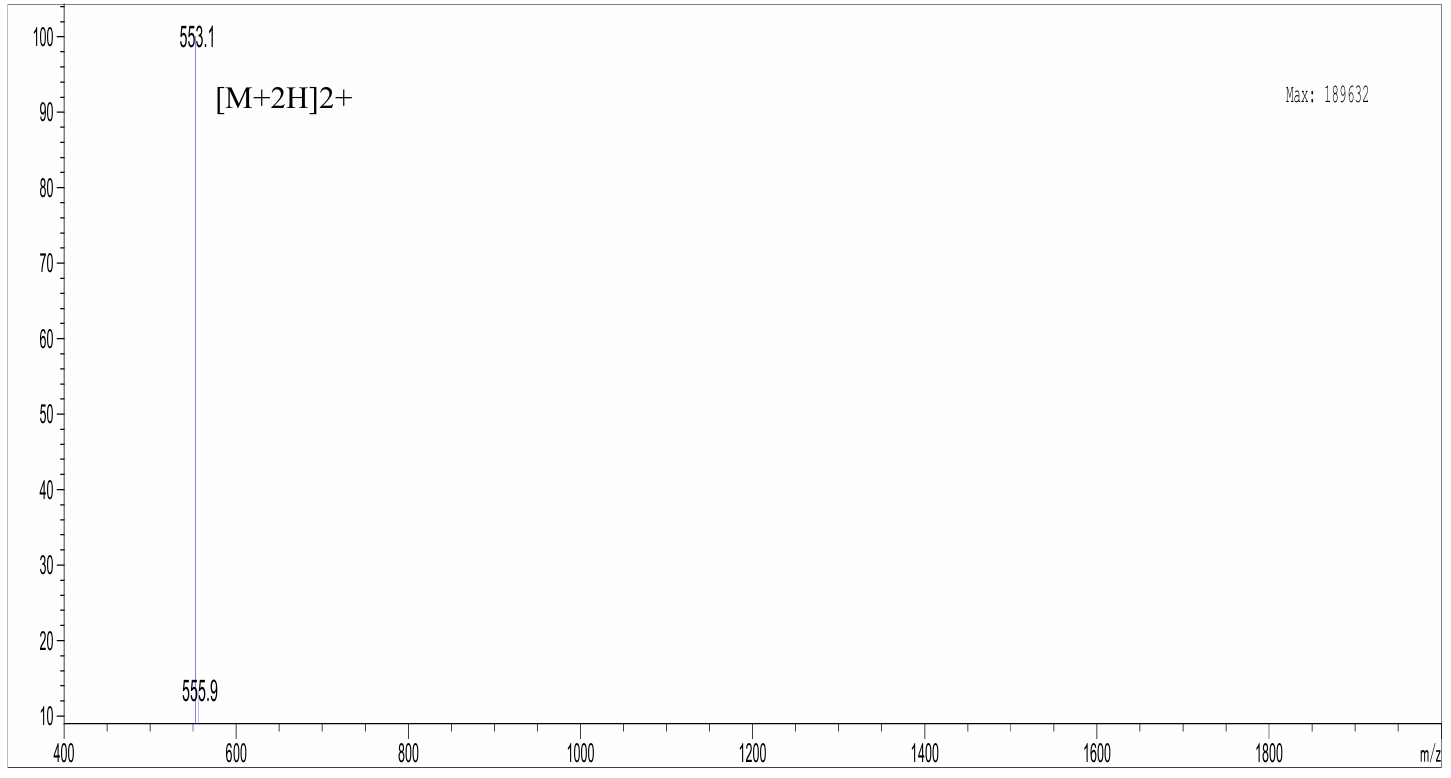

**Figure S40.** LC-MS data for Cyclic peptide 18.

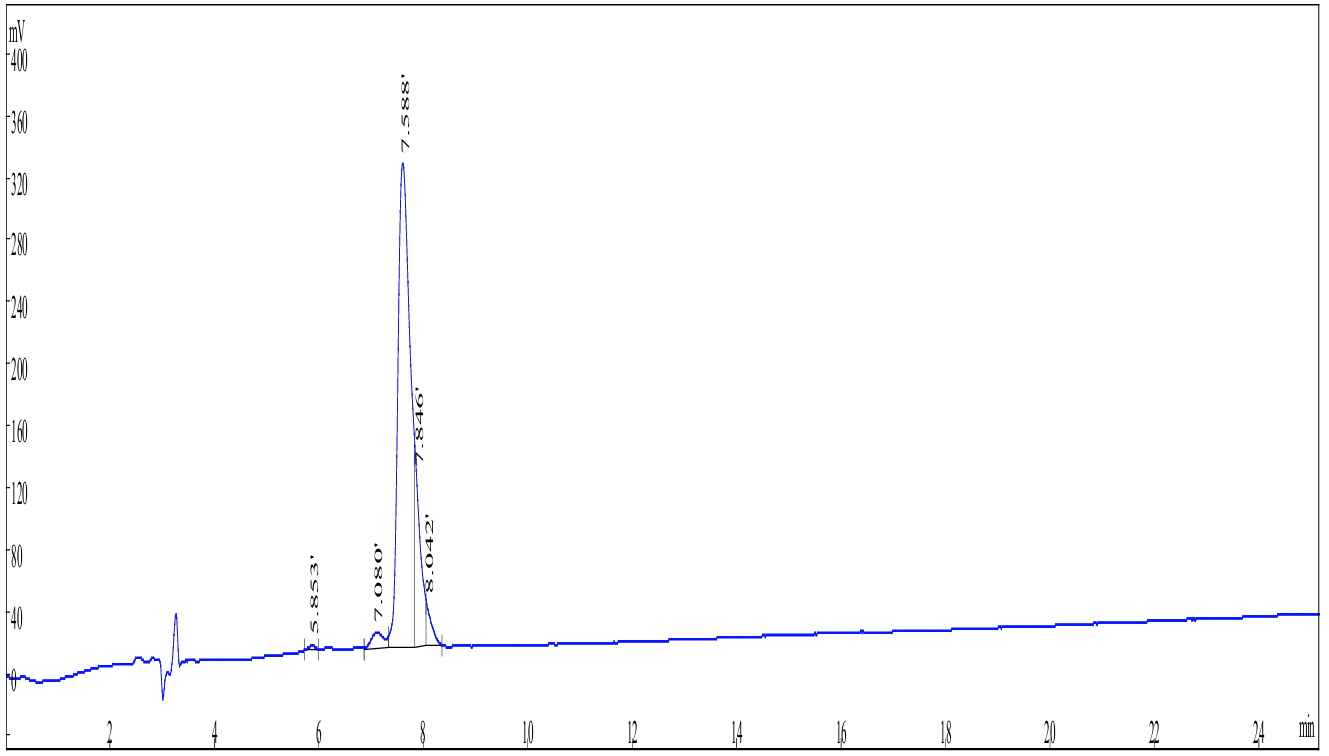

**Figure S41.** HPLC trace for Cyclic peptide 19.

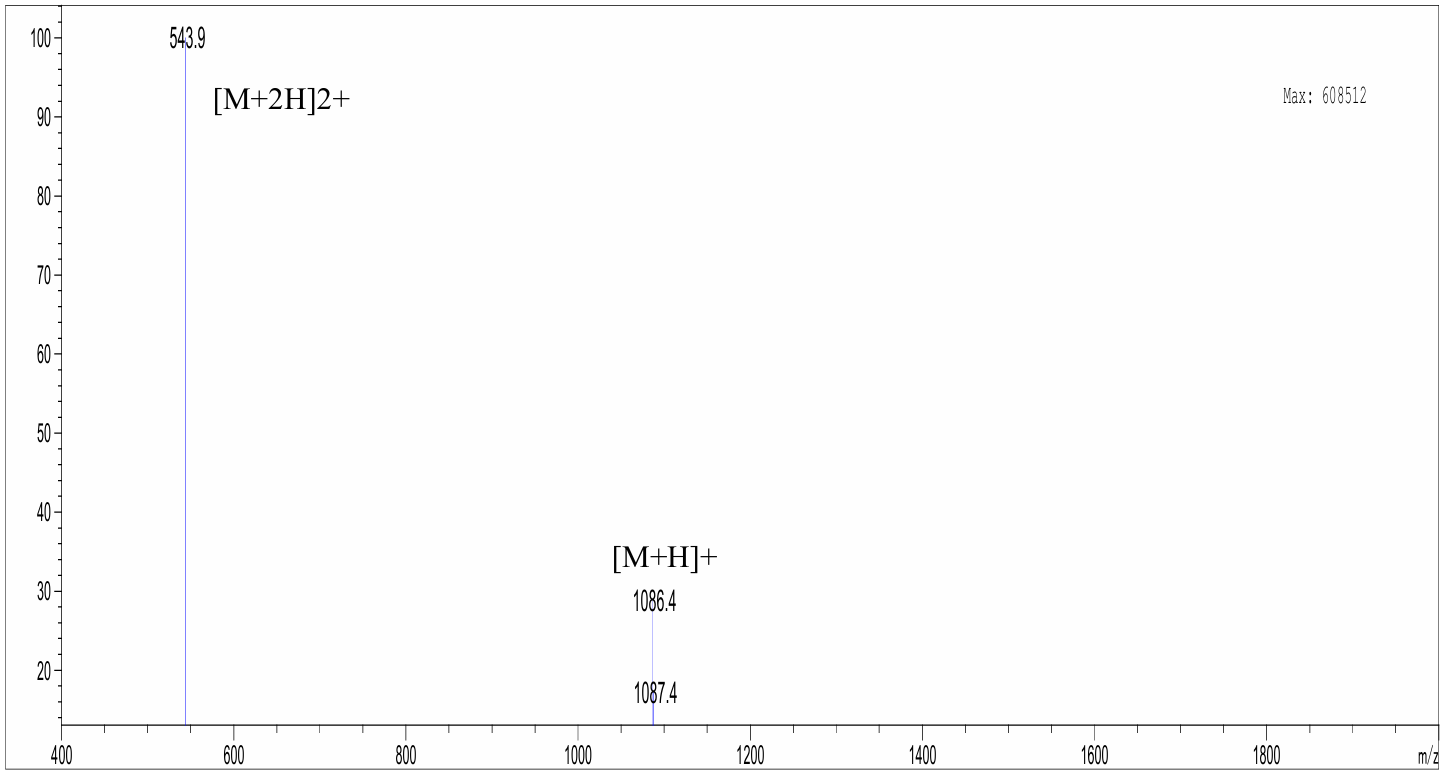

**Figure S42.** LC-MS data for Cyclic peptide 19.

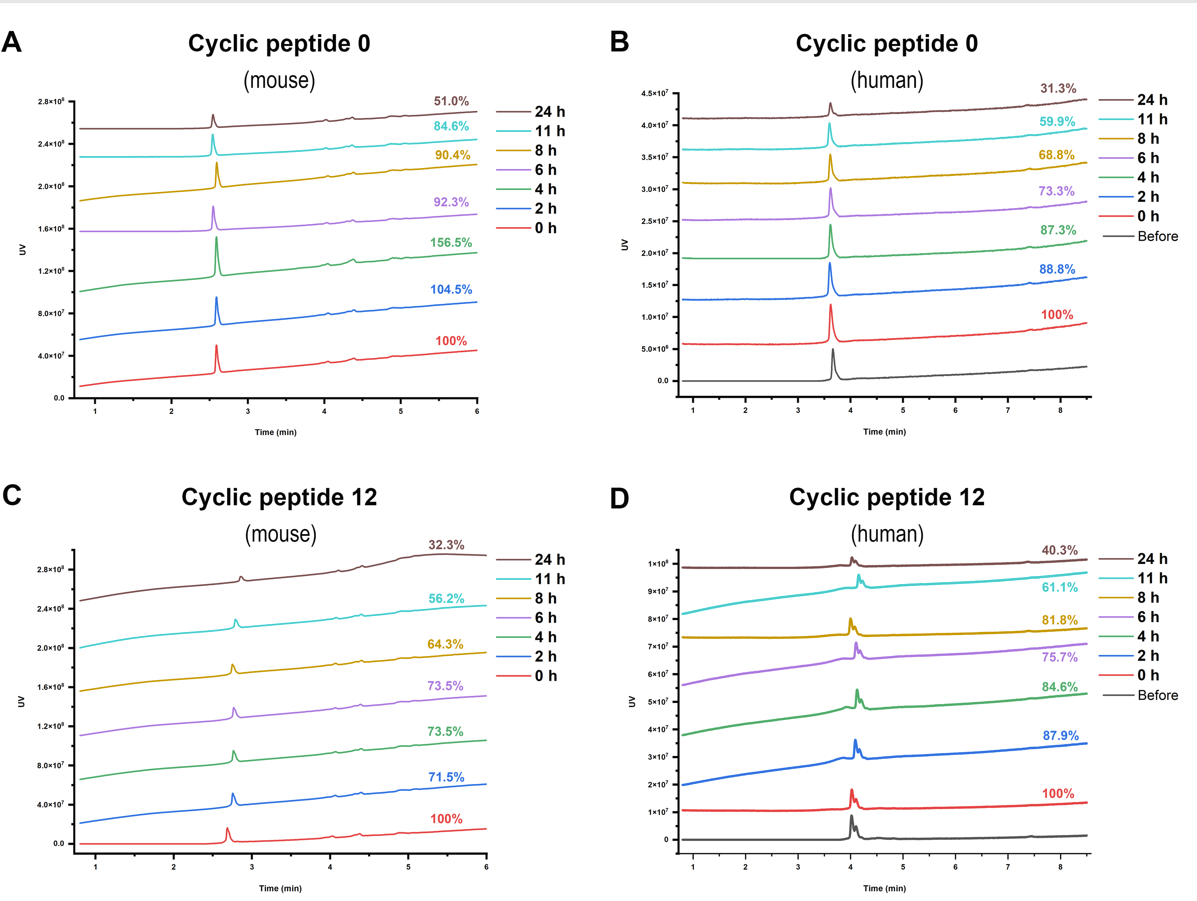

**Figure S43.** Stability of cyclic peptides in mouse and human serum. HPLC profiles showing (A, B) cyclic peptide 0 and (C, D) cyclic peptide 12 after incubation with mouse serum (left column) and human serum (right column) for varying durations. The percentages denote the remaining peptide content at each time point relative to time 0.
